# The generation of cortical novelty responses through inhibitory plasticity

**DOI:** 10.1101/2020.11.30.403840

**Authors:** Auguste Schulz, Christoph Miehl, Michael J. Berry, Julijana Gjorgjieva

**Affiliations:** Max Planck Institute for Brain Research, Frankfurt am Main, Germany; Technical University of Munich, Department of Electrical and Computer Engineering, Munich, Germany; Technical University of Munich, School of Life Sciences, Freising, Germany; Princeton University, Princeton Neuroscience Institute, Princeton, United States of America

**Author notes:** These authors contributed equally to this work (Auguste Schulz, Christoph Miehl).

## Abstract

Animals depend on fast and reliable detection of novel stimuli in their environment. Indeed, neurons in multiple sensory areas respond more strongly to novel in comparison to familiar stimuli. Yet, it remains unclear which circuit, cellular and synaptic mechanisms underlie those responses. Here, we show that inhibitory synaptic plasticity readily generates novelty responses in a recurrent spiking network model. Inhibitory plasticity increases the inhibition onto excitatory neurons tuned to familiar stimuli, while inhibition for novel stimuli remains low, leading to a network novelty response. Generated novelty responses do not depend on the exact temporal structure but rather on the distribution of presented stimuli. By including tuning of inhibitory neurons, the network further captures stimulus-specific adaptation. Finally, we suggest that disinhibition can control the amplification of novelty responses. Therefore, inhibitory plasticity provides a flexible, biologically-plausible mechanism to detect the novelty of bottom-up stimuli, enabling us to make numerous experimentally testable predictions.

## Introduction

In an ever-changing environment, animals must rapidly extract behaviorally useful information from sensory stimuli. Appropriate behavioral adjustments to unexpected changes in stimulus statistics are fundamental for the survival of an animal. We still do not fully understand how the brain detects such changes reliably and quickly. Local neural circuits perform computations on incoming sensory stimuli in an efficient manner by maximizing transmitted information or minimizing metabolic cost (Simoncelli and Olshausen, 2001; Barlow, 2013). Repeated or predictable stimuli do not provide new meaningful information. As a consequence, one should expect that responses to repeated stimuli are suppressed - a phenomenon postulated by the framework of predictive coding (Clark, 2013; Spratling, 2017). Recent experiments have demonstrated that sensory circuits across different modalities can encode a sequence or expectation violation and can detect novelty (Keller et al., 2012; Natan et al., 2015; Zmarz and Keller, 2016; Hamm and Yuste, 2016; Homann et al., 2017). The underlying neuronal and circuit mechanisms behind expectation violation and novelty detection, however, remain elusive.

A prominent paradigm used experimentally involves two types of stimuli, the repeated (or frequent) and the novel (or deviant) stimulus (Näätänen et al., 1982; Fairhall, 2014; Natan et al., 2015; Homann et al., 2017; Weber et al., 2019). Here, the neuronal responses to repeated stimuli decrease, a phenomenon that is often referred to as adaptation (Fairhall, 2014). Adaptation can occur over a wide range of timescales, which range from milliseconds to seconds (Ulanovsky et al., 2004; Lundstrom et al., 2010), and to multiple days in the case of behavioral habituation (Haak et al., 2014; Ramaswami, 2014). Responses to repeated versus novel stimuli, more generally, have also been studied on different spatial scales spanning the single neuron level, cortical microcircuits and whole brain regions. At the scale of whole brain regions, a widely studied phenomenon is the mismatch negativity (MMN), which is classically detected in electroencephalography (EEG) data and often based on an auditory or visual ‘oddball’ paradigm (Näätänen et al., 1982, 2007). The occasional presentation of the so-called oddball stimulus among frequently repeated stimuli leads to a negative deflection in the EEG signal - the MMN (Näätänen et al., 2007).

Experiments at the cellular level typically follow the oddball paradigm with two stimuli that, if presented in isolation, would drive a neuron equally strongly. However, when one stimulus is presented frequently and the other rarely, the deviant produces a stronger response relative to the frequent stimulus (Ulanovsky et al., 2003, 2004; Nelken, 2014; Natan et al., 2015). The observed reduction in response to the repeated stimulus has been termed stimulus-specific adaptation (SSA), and has been suggested to contribute to the MMN (Ulanovsky et al., 2003). SSA has been observed in multiple brain areas, most commonly reported in the primary auditory cortex (Ulanovsky et al., 2003; Yaron et al., 2012; Natan et al., 2015, 2017) and the primary visual cortex (Movshon and Lennie, 1979; Hamm and Yuste, 2016; Vinken et al., 2017; Homann et al., 2017). Along the visual pathway, SSA has also been found at different earlier stages including the retina (Schwartz et al., 2007; Geffen et al., 2007; Schwartz and Berry II, 2008) and the visual thalamic nuclei (Dhruv and Carandini, 2014; King et al., 2016).

To unravel the link between multiple spatial and temporal scales of adaptation, a variety of mechanisms has been proposed. Most notably, modeling studies have explored the role of adaptive currents, which reduce the excitability of the neuron (Brette and Gerstner, 2005) and short-term depression of excitatory feedforward synapses (Tsodyks et al., 1998). Most models of SSA in primary sensory areas of the cortex focus on short-term plasticity and the depression of thalamocortical feedforward synapses (Mill et al., 2011a,b; Park and Geffen, 2020). The contribution of other mechanisms has been under-explored in this context. Recent experimental studies suggest that inhibition and the plasticity of inhibitory synapses shape the responses to repeated and novel stimuli (Chen et al., 2015; Kato et al., 2015; Natan et al., 2015; Hamm and Yuste, 2016; Natan et al., 2017; Heintz et al., 2020). Natan and colleagues observed that in the mouse auditory cortex, both parvalbumin-positive (PV) and somatostatin-positive (SOM) interneurons contribute to SSA (Natan et al., 2015). Furthermore, neurons that are more strongly adapted receive stronger inhibitory input than less adapted neurons, suggesting potentiation of inhibitory synapses as an underlying mechanism (Natan et al., 2017). In the context of habituation, inhibitory plasticity has been previously hypothesized to be the driving mechanism to reduce neural responses to repeated stimuli (Ramaswami, 2014; Barron et al., 2017). Habituated behavior in *Drosophila*, for example, results from prolonged activation of an odor-specific excitatory subnetwork, which leads to the selective strengthening of inhibitory synapses onto the excitatory subnetwork (Das et al., 2011; Glanzman, 2011; Ramaswami, 2014; Barron et al., 2017).

Here, we focus on the role of inhibitory plasticity in characterizing neuronal responses to repeated and novel stimuli at the circuit level. We base our study on a recurrent spiking neural network model of the mammalian cortex with biologically inspired plasticity mechanisms that can generate assemblies in connectivity and attractors in activity to represent the stimulus-specific activation of specific sub-circuits (Litwin-Kumar and Doiron, 2014; Zenke et al., 2015). We model excitatory and inhibitory neurons and include stimulus-specific input not only to the excitatory but also to the inhibitory population, as found experimentally (Ma et al., 2010; Griffen and Maffei, 2014; Znamenskiy et al., 2018). This additional assumption readily leads to the formation of specific inhibitory to excitatory connections through inhibitory plasticity (Vogels et al., 2011), as suggested by recent experiments (Lee et al., 2014; Xue et al., 2014; Znamenskiy et al., 2018; Najafi et al., 2020).

We demonstrate that this model network can generate excess population activity when novel stimuli are presented as violations of repeated stimulus sequences. Our framework identifies plasticity of inhibitory synapses as a sufficient mechanism to explain population novelty responses and adaptive phenomena on multiple timescales. In addition, stimulus-specific inhibitory connectivity supports adaptation to specific stimuli (SSA). This finding reveals that the network configuration encompasses computational capabilities beyond those of intrinsic adaptation. Furthermore, we suggest disinhibition to be a powerful regulator of the amplification of novelty responses. Our modeling framework enables us to formulate additional experimentally testable predictions. Most intriguing, we hypothesize that neurons in primary sensory cortex may not signal the violation of the exact temporal sequence based on bottom-up input, but rather adapt to the distribution of presented stimuli.

## Results

### A recurrent neural network model with plastic inhibition can generate novelty responses

Recent experimental studies have indicated an essential role of inhibitory circuits and inhibitory plasticity in adaptive phenomena and novelty responses (Chen et al., 2015; Kato et al., 2015; Natan et al., 2015; Hamm and Yuste, 2016; Natan et al., 2017; Heintz et al., 2020). To understand if and how plastic inhibitory circuits could explain the emergence of novelty responses, we built a biologically-plausible spiking neuronal network model of recurrently connected excitatory and inhibitory neurons based on recent experimental findings on tuning, connectivity, and inhibitory and excitatory spike-timing-dependent synaptic plasticity (STDP) in the cortex (Methods) (Pfister and Gerstner, 2006; Vogels et al., 2011; Fiete et al., 2010; Sjöström et al., 2001; D’amour and Froemke, 2015). We targeted different subsets of excitatory and inhibitory neurons with different external stimuli, to model that these neurons are stimulus-specific (‘tuned’) to a given stimulus (Fig. 1A, left, see Methods). One neuron can be driven by multiple stimuli. Starting from an initially randomly connected network, presenting tuned input led to the emergence of excitatory assemblies, which are strongly connected, functionally related subsets of excitatory neurons (Suppl. Fig. S1C, left). Furthermore, tuned input also led to the stimulus-specific potentiation of inhibitory to excitatory connections (Suppl. Fig. S1E, left). We refer to this part of structure formation as the ‘pretraining phase’ of our simulations (Methods). This pretraining phase imprints structure in the network prior to the actual stimulation paradigm and could correspond to the activity-dependent refinement of structured connectivity during early postnatal development (Thompson et al., 2017).

**Figure 1.**
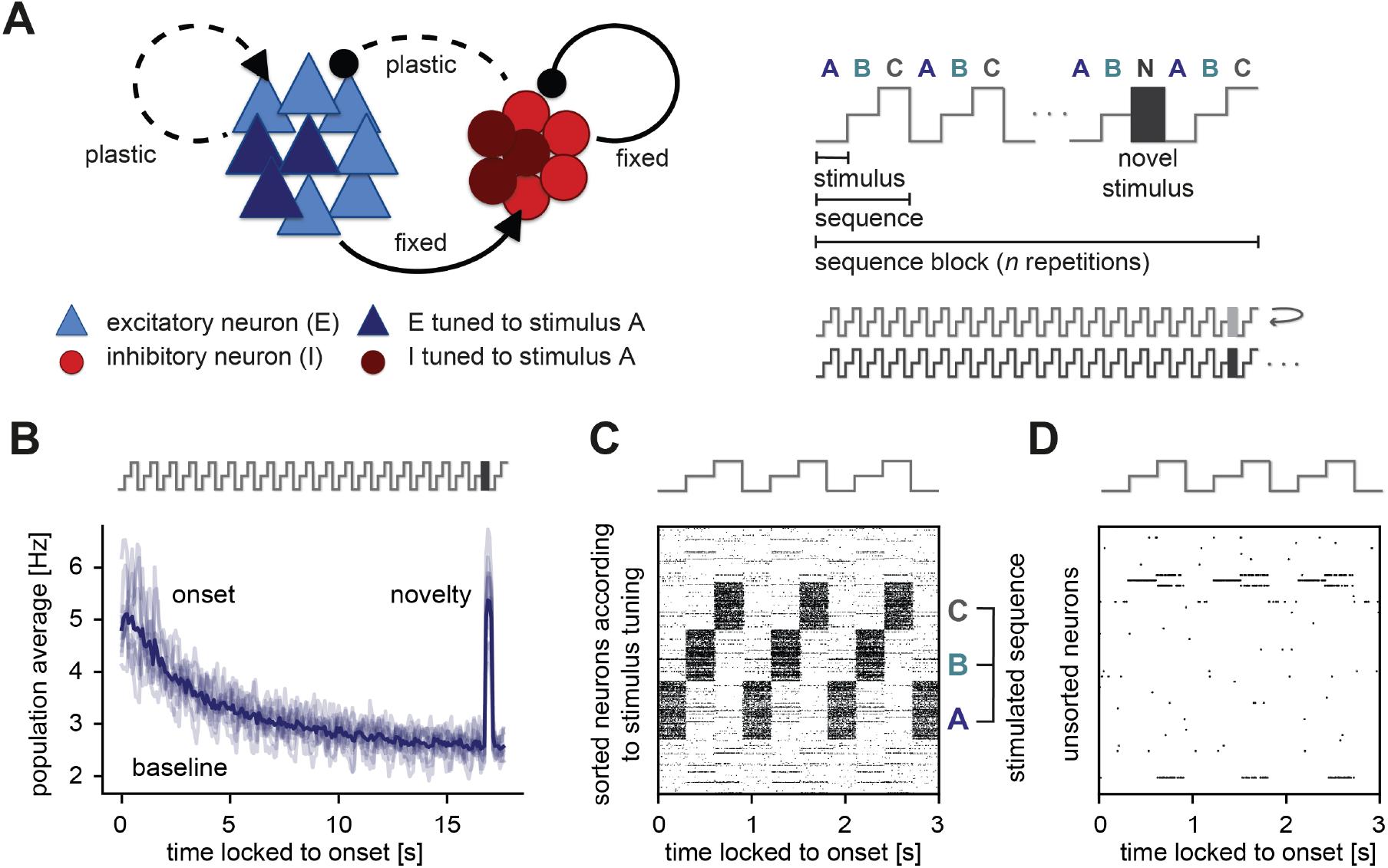
Generation of novelty responses in a recurrent plastic neural network model. **A.** Left: A recurrently connected network of excitatory (E) neurons (blue triangles) and inhibitory (I) neurons (red circles) receiving tuned input. Excitatory neurons tuned to a sample stimulus A are highlighted in dark blue, the inhibitory counterparts in dark red. E-to-E synapses and I-to-E synapses were plastic, and all other synapses were fixed. Right: Schematic of the stimulation protocol. Multiple stimuli (A, B, and C) were presented in a sequence (ABC). Each sequence was repeated *n* times in a sequence block. In the second-to-last sequence, the last stimulus was replaced by a novel stimulus (N). Multiple sequence blocks followed each other without interruption, with each block containing sequences of different stimuli. **B.** Population average firing rate of all excitatory neurons as a function of time after the onset of a sequence block. Activity was averaged (solid line) across multiple non-repeated sequence blocks (transparent lines: individual blocks). A novel stimulus (dark gray) was presented as the last stimulus of the second-to-last sequence. **C.** Spiking activity in a subset of 1000 excitatory neurons where the neurons were sorted according to the stimulus from which they receive tuned input. A neuron can receive input from multiple stimuli and can show up more than once in this raster plot. Time was locked to the sequence block onset. **D.** A random unsorted subset of 50 excitatory neurons.

To test the influence of inhibitory plasticity on the emergence of a novelty response, we followed an experimental paradigm used to study novelty responses in layer 2/3 (L2/3) of mouse primary visual cortex (V1) (Homann et al., 2017) (Fig. 1A, right). Three different stimuli (A, B, and C) were presented in a sequence (ABC). The same sequence (ABC) was then repeated several times in a sequence block. In the second-to-last sequence, the last stimulus was replaced by a novel stimulus (N). In the consecutive sequence block, a new sequence with different stimuli was presented (we referred to this as a ‘unique sequence block experiment’). The novel stimuli were also different for each sequence block. In this paradigm, we observed elevated population activity in the excitatory population at the beginning of each sequence block (‘onset response’) and a steady reduction to a baseline activity level for the repeated sequence presentation (Fig. 1B). Upon presenting a novel stimulus, the excitatory population showed excess activity, clearly discernible from baseline, called the’novelty response’. This novelty response was comparable in strength to the onset response. Sorting spike rasters according to sequence stimuli revealed that stimulation leads to high firing rates in the neurons that are selective to the presented stimulus (A, B, or C) (Fig. 1C). When examining a random subset of neurons, we found general response sparseness and periodicity during sequence repetitions (Fig. 1D), very similar to experimental findings (Homann et al., 2017). More concretely, sparse population activity for repeated stimuli in our model network was the result of each stimulus presentation activating a subset of excitatory neurons in the network, which were balanced by strong inhibitory feedback. Therefore, only neurons that directly received this feedforward drive were highly active, while most other neurons in the network were instead rather silent. Periodicity in the activity of single neurons resulted from the repetition of a sequence.

Our results suggest that presenting repeated stimuli (and repeated sequences of stimuli) to a plastic recurrent network with tuned excitatory and inhibitory neurons readily leads to a reduction of the excitatory averaged population response, consistent with the observed adaptation in multiple experimental studies in various animal models and brain regions (Ulanovsky et al., 2003; Hamm and Yuste, 2016; Homann et al., 2017). Importantly, the model network generates a novelty response when presenting a novel stimulus by increasing the excitatory population firing rate at the time of stimulus presentation (Näätänen et al., 2007).

### The dynamics of novelty and onset responses depend on sequence properties

To explore the dynamics of novelty responses, we probed the model network with a modified stimulation paradigm. Rather than fixing the number of sequence repetitions in one sequence block (Fig. 1A, right), here we presented a random number of sequence repetitions (nine values between 4 and 45 repetitions) for each sequence block. This allowed us to measure the novelty and onset responses as a function of the number of sequence repetitions. Novelty and onset responses were observed after as few as four sequence repetitions (Fig. 2A). After more than 15 sequence repetitions, the averaged excitatory population activity reached a clear baseline activity level (Fig. 2A). The novelty response amplitude, measured by the population rate of the novelty peak minus the baseline population rate, increased with the number of sequence repetitions before saturating for a high number of sequence repeats (Fig. 2B, black dots). The onset response amplitude after the respective sequence block followed the same trend (Fig. 2B, grey dots). Next, we varied the number of stimuli in a sequence, resulting in different sequence lengths across blocks (3 to 15 stimuli per sequence). By averaging excitatory population responses across sequence blocks with equal length, we found that the decay of the onset response depends on the number of stimuli in a sequence (Fig. 2C). Upon fitting an exponentially decaying function to the activity of the onset response, we derived a linear relationship between the number of stimuli in a sequence and the decay constant (Fig. 2D).

**Figure 2.**
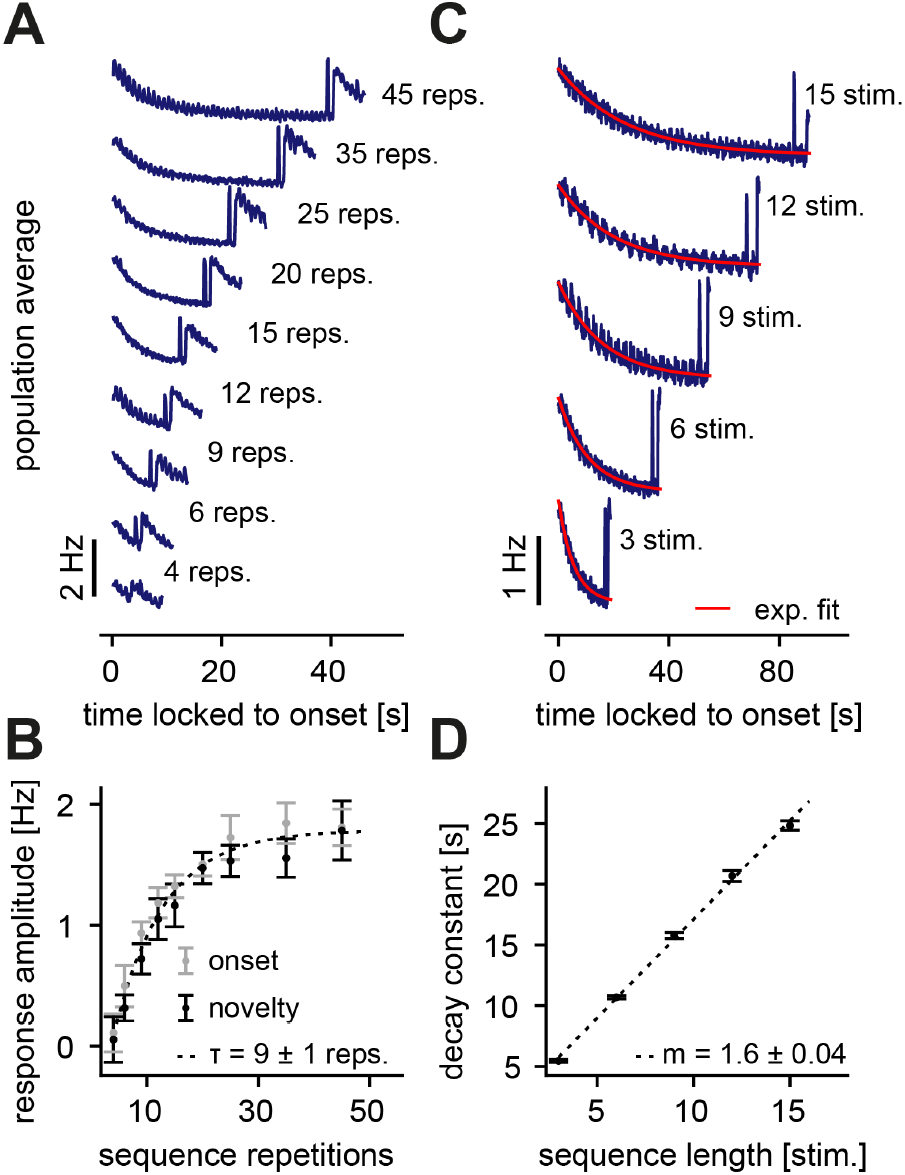
Dependence of the novelty response on the number of sequence repetitions and the sequence length. **A.** Population average firing rate for a different number of sequence repetitions within a sequence block. Time is locked to the sequence block onset. **B.** The response amplitude of the onset (gray) and the novelty (black) response as a function of sequence repetitions fit with an exponential with a time constant *τ*. **C.** Population average firing rate of all excitatory neurons for varying sequence length fit with an exponential function (red). Time is locked to the sequence block onset. **D.** The onset decay time constant (fit with an exponential as shown in panel C) as a function of sequence length. The simulated data is fit with a linear function with slope *m*. **B, D.** Error bars correspond to the standard deviation across five simulated instances of the model.

In summary, we found that novelty responses arise for different sequence variations. Our model network suggests that certain features of the novelty response depend on the properties of the presented sequences. Changing the number of sequence repetitions modifies the onset and novelty response amplitude (Fig. 2A, B), while a longer sequence length leads to a longer adaptation time constant (Fig. 2C, D). Interestingly, both findings are in good qualitative agreement with experimental data that presented similar sequence variations (Homann et al., 2017). An exponential fit of the experimental data found a time constant of *τ* = 3.2 ± 0.7 repetitions when the number of sequence repetitions was varied (Homann et al., 2017). The time constant in our model network was somewhat longer (*τ* = 9 ± 1 repetitions), but similar order of magnitude (Fig. 2B). Similarly, our model network produced a linear relationship between the adaptation time constant and sequence length with a slope of *m* = 1.6 ± 0.04 (Fig. 2D), very close to the slope extracted from the data (*m* = 2.1 ± 0.3) (Homann et al., 2017). Therefore, grounded on biologically-plausible plasticity mechanisms, and capable of capturing the emergence and dynamics of novelty responses, our model network provides a suitable framework for a mechanistic dissection of the circuit contributions in the generation of a novelty response.

### The novelty response is independent of a sequence’s temporal structure

Experimental studies have often reported novelty or deviant responses by averaging across several trials due to poor signal to noise ratios of the measured physiological activity (Homann et al., 2017; Vinken et al., 2017). Therefore, we investigated the network response to paradigms with repeated individual sequence blocks (Fig. 3A). We randomized the order of the sequence block presentation to avoid additional temporal structure beyond the stimulus composition of the sequences. Repeating sequence blocks dampened the onset response at sequence onset compared to the unique sequence block conditions (compare Fig. 1B and Fig. 2A,B with Fig. 3A). Next, we wondered whether the excitatory and inhibitory population responses to repeated and novel stimuli are related. We found that both excitatory and inhibitory populations adapt to the repeated stimuli and show a prominent novelty peak that is larger than the respective averaged onset response (Fig. 3B,C). Based on these findings, we make the following predictions for future experiments: (1) A novelty response is detectable in both the excitatory and inhibitory populations. (2) The sequence onset response is dampened for multiple presentations of the same sequence block compared to the presentation of unique sequence blocks.

**Figure 3.**
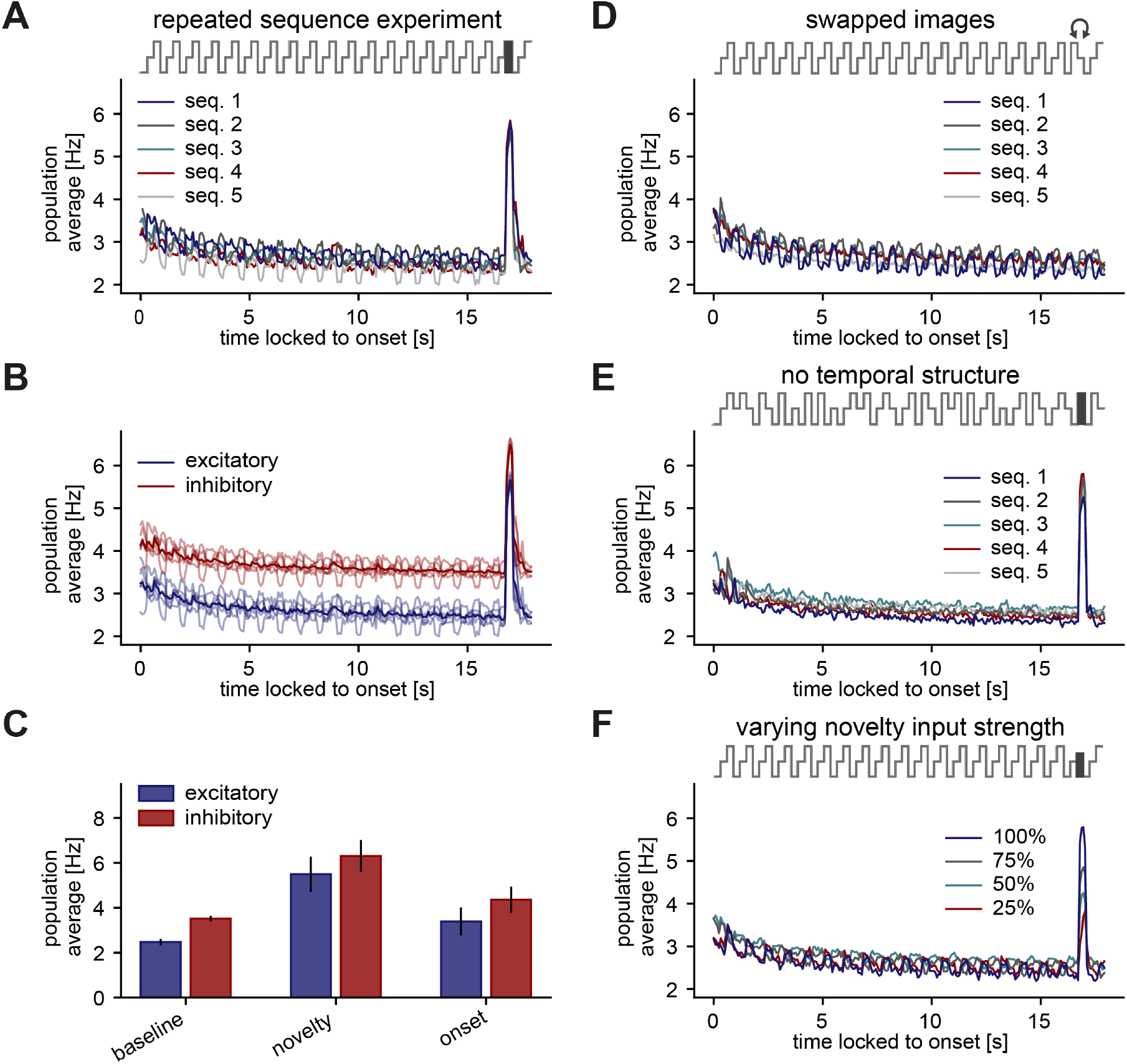
Neurons do not learn the exact temporal structure of a sequence. **A.-F.** Population average firing rate of all excitatory neurons (and all inhibitory neurons in B,C) during the presentation of five different repeated sequence blocks. The population firing rate is averaged across ten repetitions of each sequence block. Time is locked to sequence block onset. **A.** A novel stimulus is presented as the last stimulus of the second-to-last sequence. **B.** Same as panel A but for both excitatory and inhibitory populations (transparent lines: individual sequence averages). **C.** Comparison of baseline, novelty, and onset response for inhibitory and excitatory populations. Error bars correspond to the standard deviation across the five sequence block averages shown in **B. D.** In the second-to-last sequence, the last and second-to-last stimulus are swapped instead of presenting a novel stimulus. **E.** Within a sequence, stimuli are shuffled in a pseudo-random manner where a stimulus cannot be presented twice in a row. A novel stimulus is presented as the last stimulus of the second-to-last sequence. **F.** A novel stimulus is presented as the last stimulus of the second-to-last sequence. Each sequence has a different feedforward input drive for the novel stimulus, indicated by the percentage of the typical input drive for the novel stimulus used before.

Next, we investigated whether the novelty responses observed in the model network depend on the temporal structure of the sequence. If the novelty responses were to truly signal the violation of temporal structure or the stimulus predictability in a sequence, we would expect a novelty response to occur if two stimuli in a sequence were swapped, i.e. ACB instead of ABC. We found that swapping the last and second-to-last stimulus, instead of presenting a novel stimulus, does not elicit a novelty response (Fig. 3D). Additionally, we asked whether the exact temporal structure of the stimuli within a sequence influences the novelty response. Shuffling the stimuli within a sequence block generated the same novelty response and adaptation to the repeated stimuli as in the strictly sequential case (Fig. 3E, compare to A). Finally, we investigated if the novelty peak depends on the input firing rate of the novel stimulus. We found that a reduction of the input drive indeed decreases the novelty peak, revealing a monotonic dependence of the novelty response on stimulus strength (Fig. 3F). Based on these results, we make two additional predictions: (3) A novelty response is independent of the exact temporal structure of the sequences, hence it encodes the distribution of presented stimuli, rather than the temporal structure of a sequence. (4) A novelty response depends on the strength of the novel stimulus.

### Increased inhibition onto highly active neurons leads to adaptation

To gain an intuitive understanding for the sensitivity of novelty responses to stimulus identity but lack of sensitivity to temporal structure, we more closely examined the role of inhibitory plasticity as the leading mechanism behind the novelty responses in our model. We found that novelty responses arise because inhibitory plasticity fails to sufficiently increase inhibitory input and to counteract the excess excitatory input into excitatory neurons upon the presentation of a novel stimulus. In short, novelty responses can be understood as the absence of adaptation in an otherwise adapted response. Adaptation in the network arises through increased inhibition onto highly active neurons through selective strengthening of I-to-E weights (Fig. 4A).

**Figure 4.**
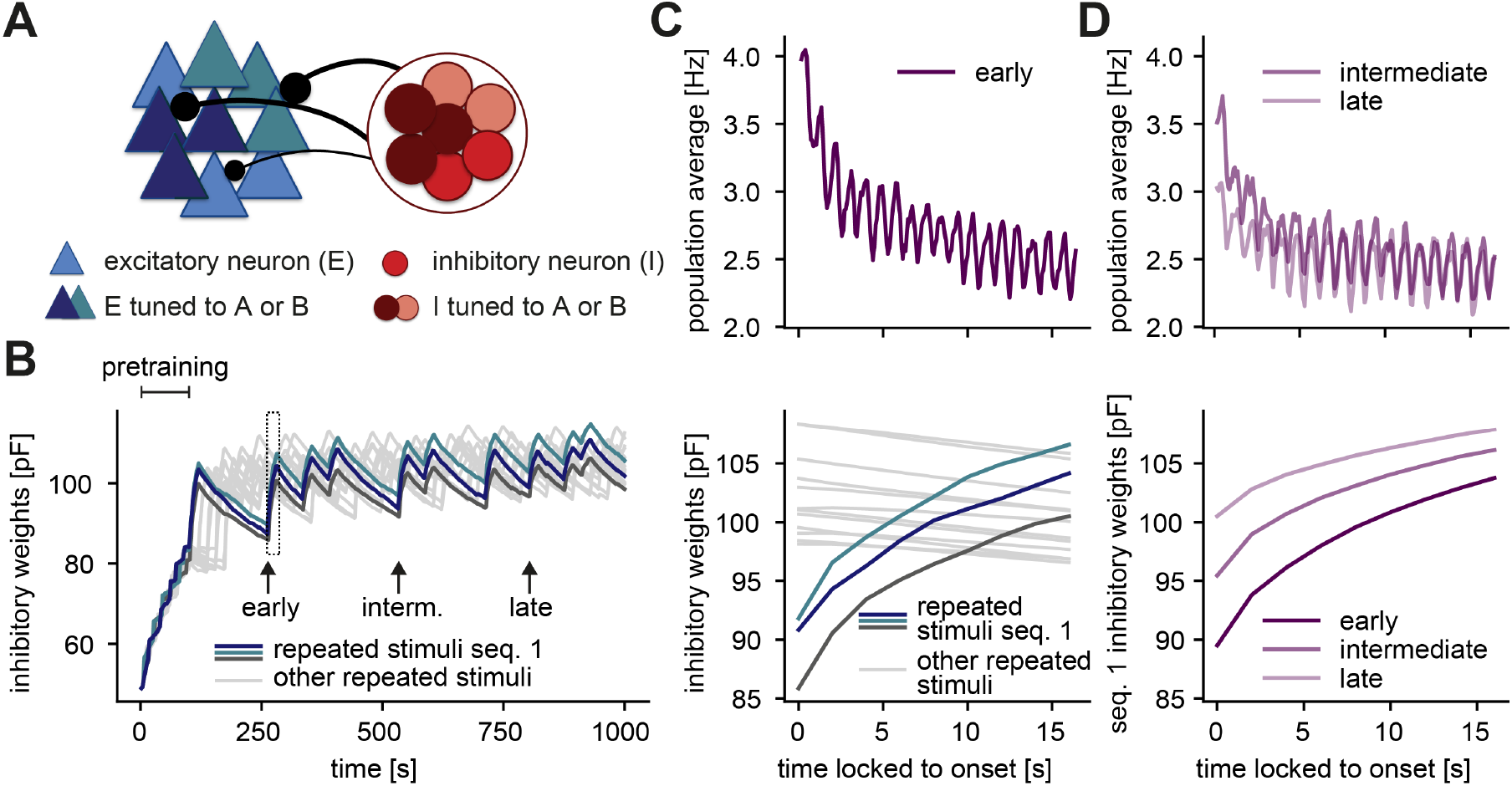
Inhibition onto neurons tuned to repeated stimuli increases during sequence repetitions. **A.** Schematic of increased inhibitory weights onto two stimulus-specific assemblies upon the repeated presentation of stimuli A and B (indicated in dark blue and turquoise) relative to neurons from other assemblies (light blue). **B.** Evolution of the average inhibitory weights onto stimulus-specific assemblies. Colored traces mark three stimulus-specific assemblies in sequence 1: A, B, and C. Arrows indicate time points of early, intermediate, and late sequence block presentation shown in C and D. **C.** Top: Population average firing rate of all excitatory neurons during the repeated presentation of sequence 1 at an early time point (see panel B). Time is locked to sequence onset. Bottom: Close-up of panel B (rectangle). Time is locked to sequence onset. **D.** Top: Same as panel C (top) but at intermediate and late time points (see panel B). Bottom: Corresponding dynamics of the average inhibitory weights onto all three stimulus-specific assemblies from sequence 1 at early, intermediate and late time points (see panel B). The dark purple trace (early) corresponds to the average of the three colored traces in C (bottom).

To determine how inhibitory plasticity drives the generation of novelty responses or, equivalently, adaptation in our model, we studied the evolution of inhibitory weights. The inhibitory weights onto stimulus-specific assemblies tuned to the stimuli in a given sequence increased upon presentation of the corresponding sequence block, and decreased otherwise (Fig. 4B). The population firing rate during repeated presentation of a sequence decreased (adapted) on the same timescale as the increase of the inhibitory weights related to this sequence (Fig. 4C). When a stimulus was presented to the network for the first time, the total excitatory input to the corresponding excitatory neurons was initially not balanced by inhibition. Hence, the neurons within the assembly tuned to that stimulus exhibited elevated activity at sequence onset, leading to what we called the‘onset response’(Fig. 1B). The same was true for the novelty responses as reflected in low inhibitory weights onto novelty assemblies relative to repeated assemblies (Suppl. Fig. S1D). Consequently, the generation of a novelty response was independent on the specific temporal order of the stimuli within a sequence (Fig. 3). Swapping two stimuli did not generate a novelty response since the corresponding assemblies of each stimulus were already in an adapted state. Therefore, we conclude that the exact temporal structure of stimulus presentations was not relevant for the novelty response, as long as the overall distribution of stimuli was maintained.

Interestingly, we found that adaptation occurs on multiple timescales in our model. The fastest is the timescale of milliseconds on which inhibitory plasticity operates, the next slowest is the timescale of seconds corresponding to the presentation of a sequence block, and finally the slowest is the timescale of minutes corresponding to the presentation of the same sequence block multiple times (Fig. 4D, top; also compare Fig. 1B and Fig. 3A). The slowest decrease in the population firing rate results from long-lasting changes in the average inhibitory weights onto the excitatory neurons tuned to the stimuli within a given sequence. Hence, the average inhibitory weight for a given sequence increases with the number of previous sequence block presentations of that sequence (Fig. 4D, bottom).

In summary, we identified the plasticity of connections from inhibitory to excitatory neurons belonging to a stimulus-specific assembly as the key mechanism in our framework for the generation of novelty responses and for the resulting adaptation of the network response to repeated stimuli. This adaptation occurs on multiple timescales, covering the range from the timescale of inhibitory plasticity (milliseconds) to sequence block adaptation (seconds) to the presentation of multiple sequence blocks (minutes).

### Inhibitory plasticity and tuned inhibitory neurons support stimulus-specific adaptation

Next, we investigated whether inhibitory plasticity of tuned inhibitory neurons support additional computational capabilities beyond the generation of novelty responses and adaptation of responses to repeated stimuli on multiple timescales. Therefore, we implemented a different stimulation paradigm, known as stimulus-specific adaptation (SSA). Atthe single cell level, SSA typically involves a so-called oddball paradigm where two stimuli elicit equally strong response when presented in isolation, but when one is presented more frequently, the elicited response is weaker than for a rarely presented stimulus (Natan et al., 2015).

We implemented a similar paradigm at the network level where the excitatory neurons corresponding to two stimuli A and B were completely overlapping and the inhibitory neurons were partially overlapping (Fig. 5A). Upon presenting stimulus A several times, the neuronal response gradually adapted to the baseline level of activity, while presenting the oddball stimulus B resulted in an increased population response (Fig. 5B). Therefore, this network was able to generate SSA. Even though stimuli A and B targeted the same excitatory cells, the network response adapted only to stimulus A, while generating a novelty response for stimulus B. Even after presenting stimulus B, activating stimulus A again preserved the adapted response (Fig. 5B). This form of SSA exhibited by our model network is in agreement with many experimental findings in the primary auditory cortex, primary visual cortex, and multiple other brain areas and animal models (Nelken, 2014). In our model network, SSA could neither be generated with adaptive neurons and static synapses (Fig. 5C, top;Methods), nor with inhibitory plasticity without inhibitory tuning (Fig. 5C, bottom). In fact, including an adaptive current in the constituent neurons of the network neurons (Brette and Gerstner, 2005) did not even lead to adaptation of the response to a frequent stimulus since firing rates rapidly adapted during stimulus presentation and completely recovered in the inter-stimulus pause (Fig. 5C, top).

**Figure 5.**
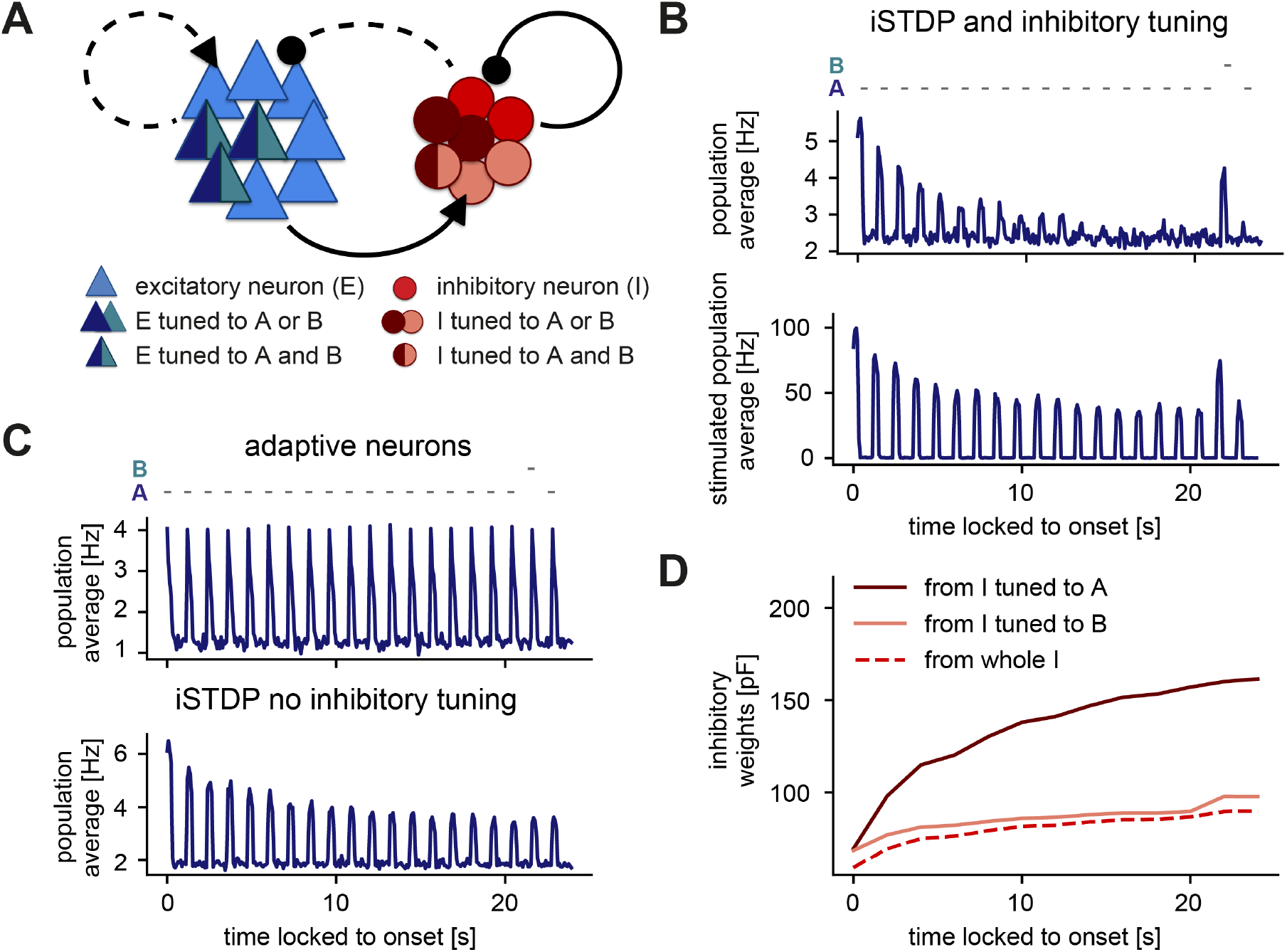
Stimulus-specific adaptation follows from inhibitory plasticity and tuning of both excitatory and inhibitory neurons. **A.** Stimuli A and B drive the same excitatory neurons (dark blue and turquoise). Some neurons in the inhibitory population are driven by both A and B (dark red and rose) and some are driven by only one of the two stimuli (dark red or rose). **B.,C.** Population average firing rate of excitatory neurons over time while stimulus A is presented 20 times. Stimulus B is presented instead of A as the second-to-last stimulus. Time is locked to stimulation onset. **B.** Top: Population average of all excitatory neurons in the network with inhibitory plasticity (iSTDP) and inhibitory tuning. Bottom: Population average of stimulated excitatory neurons only (stimulus-specific to A and B). **C.** Top: Same as panel B (top) for neurons with an adaptive current in a non-plastic recurrent network. Bottom: Same as panel B (top) for the network with inhibitory plasticity (iSTDP) and no inhibitory tuning. **D.** Weight evolution of stimulus-specific inhibitory weights corresponding to stimuli A and B and average inhibitory weights.

We investigated the dynamics of inhibitory weights to understand the mechanism behind SSA in our model network. During the presentation of stimulus A, stimulus-specific inhibitory weights corresponding to stimulus A (average weights from inhibitory neurons tuned to stimulus A onto excitatory neurons tuned to stimulus A, see Suppl. Fig. S1A, right) increased their strength, while stimulus-specific inhibitory weights corresponding to stimulus B remained low (Fig. 5D). Hence, upon presenting the oddball stimulus B, the stimulus-specific inhibitory weights corresponding to stimulus B remained sufficiently weak to keep the firing rate of excitatory neurons high, thus resulting in a novelty response.

Therefore, our results suggest that the combination of inhibitory plasticity and inhibitory tuning can give rise to SSA. Previous work has argued that inhibition or inhibitory plasticity does not allow for SSA (Nelken, 2014). However, this is only true if inhibition is interpreted as a ‘blanket’ without any tuning in the inhibitory population. Including recent experimental evidence for tuned inhibition into the model, (Lee et al., 2014; Xue et al., 2014; Znamenskiy et al., 2018), can indeed capture the emergence of SSA.

### Disinhibition leads to novelty response amplification and a dense population response

Beyond the bottom-up computations captured by the network response to the different stimuli, we next explored the effect of additional modulations or top-down feedback into our network model. Top-down feedback has been frequently postulated to signal the detection of an error or irregularity in the framework of predictive coding (Clark, 2013; Spratling, 2017). Therefore, we specifically tested the effect of disinhibitory signals on sequence violations by inhibiting the population of inhibitory neurons during the presentation of a novel stimulus (Fig. 6A). Recent evidence has identified a differential disinhibitory effect in sensory cortex in the context of adapted and novelty responses (Natan et al., 2015). However, due to the scarcity of detailed knowledge about higher-order feedback signals or within-layer modulations in this context, we did not directly model the source of disinhibition.

**Figure 6.**
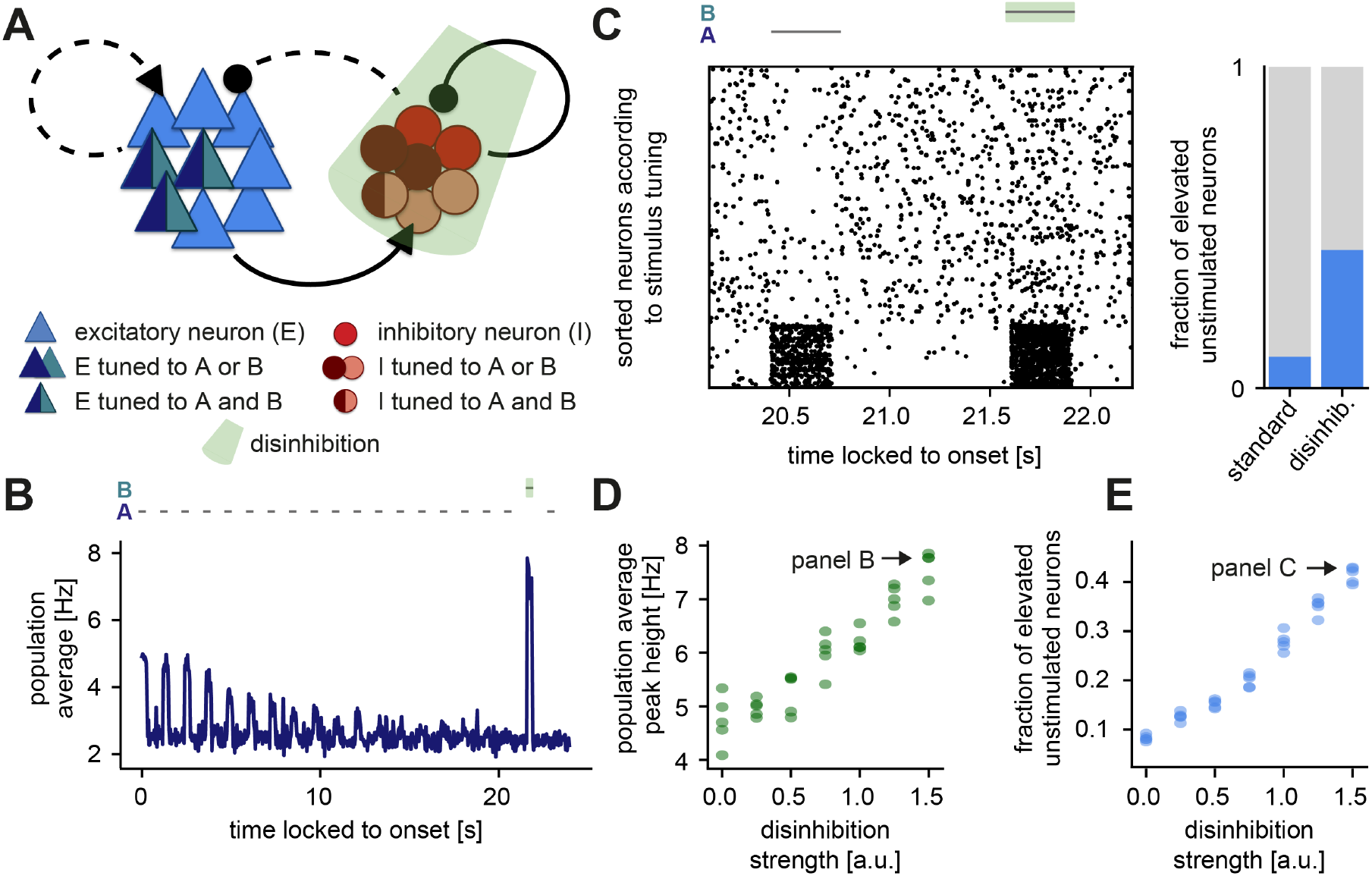
Disinhibition leads to a novelty response amplification and a dense population response. **A.** Stimuli A and B drive the same excitatory neurons (dark blue and turquoise). Some neurons in the inhibitory population are driven by both A and B (dark red and rose) and some are driven by only one of the two stimuli (dark red or rose). Inhibition (light green) of the entire inhibitory population leads to disinhibition of the excitatory population. **B.** Population average firing rate of all excitatory neurons over time while stimulus A is presented 20 times. Stimulus B is presented instead of A as the second-to-last stimulus. During the presentation of B, the inhibitory population is inhibited. Time is locked to stimulation onset. **C.** Left: Raster plot of 250 excitatory neurons corresponding to the population average shown in panel B. The 50 neurons in the bottom part of the raster plot are tuned to stimuli A and B. Time is locked to stimulation onset. Right: Fraction of stimulus-unspecific (unstimulated) neurons (light blue triangles in panel A) that have a higher firing rate during the the presentation of a stimulus than in the 300 ms preceding it. The raster plot and the fraction of elevated unstimulated neurons are shown for the presentation of stimulus B and the preceding presentation of stimulus A. **D.** Response amplitude during disinhibition and the presentation of stimulus B, as a function of the disinhibition strength. Arrow indicates the response amplitude of the trace shown in panel B. Results are shown for five simulations. **E.** Fraction of elevated unstimulated neurons during disinhibition as a function of the disinhibition strength. Arrow indicates the data point corresponding to panel C. Results are shown for five simulations.

When repeating the SSA experiment (Fig. 5) and applying such a disinhibitory signal (inhibition of the inhibitory population) at the time of the novel stimulus B, our model network amplified the novelty response (Fig. 6B, shaded green, also compare to Fig. 5C, top). Disinhibition also increased the density of the network response which corresponds to the number of active excitatory neurons (Fig. 6C, left). Indeed, disinhibition increased the fraction of stimulus-unspecific (unstimulated) neurons with elevated activity, which is defined as the fraction of neurons showing more activity during the presentation of a stimulus (to which they are not tuned) than in the 300 ms preceding it (Fig. 6C, right). Dense novelty responses have been recently reported experimentally, where novel stimuli elicited excess activity in a large fraction of the neuronal population in mouse V1 (Homann et al., 2017). Given that the inclusion of disinhibitory signals readily leads to increased density, we suggest that disinhibition might underlie these experimental findings.

In sum, we found that by controlling the total disinhibitory strength (Methods), disinhibition can flexibly amplify the novelty peak (Fig. 6D) and increase the density of novelty responses (Fig. 6E). Therefore, we propose that disinhibition can be a powerful mechanism to modulate novelty responses in a network of excitatory and inhibitory neurons.

## Discussion

We developed a recurrent network model with plastic synapses to unravel the mechanistic underpinning of adaptive phenomena and novelty responses. Using the paradigm of repeated stimulus sequences (Fig. 1A, right), our model network captured the adapted, sparse and periodic responses to repeated stimuli (Fig. 1B-D) as observed experimentally (Fairhall, 2014; Homann et al., 2017). The model network also exhibited a transient elevated population responses to novel stimuli (Fig. 1B), which could be modulated by the number of sequence repetitions and the sequence length in the stimulation paradigm (Fig. 2), in good qualitative agreement with experimental data (Homann et al., 2017). We proposed inhibitory synaptic plasticity as a key mechanism behind the generation of these novelty responses. In our model, repeated stimulus presentation triggered inhibitory plasticity onto excitatory neurons selective to the repeated stimulus, reducing the response of excitatory neurons and resulting in their adaptation (Fig. 4). In contrast, for a novel stimulus inhibitory input onto excitatory neurons tuned to that stimulus remained low, generating the elevated novelty response.

Based on experimental evidence (Ohki and Reid, 2007; Griffen and Maffei, 2014), we included specific input onto both the excitatory and the inhibitory populations (Fig. 1A, left). Such tuned inhibition (as opposed to untuned, ‘blanket’ inhibition commonly used in previous models) enabled the model network to generate SSA (Fig. 5). Additionally, in the presence of tuned inhibition, a top-down disinhibitory signal achieved a flexible control of the amplitude and density of novelty responses (Fig. 6). Therefore, besides providing a mechanistic explanation for the generation of adapted and novelty responses to repeated and novel sensory stimuli, respectively, our network model enabled us to formulate multiple experimentally testable predictions, as we describe below.

### Inhibitory plasticity as an adaptive mechanism

We proposed inhibitory plasticity as the key mechanism that allows for adaptation to repeated stimulus presentation and the generation of novelty responses in our model. Many experimental studies have characterized spike-timing-dependent plasticity (STDP) of synapses from inhibitory onto excitatory neurons (Holmgren and Zilberter, 2001; Woodin et al., 2003; Haas et al., 2006; Maffei et al., 2006; Wang and Maffei, 2014; D’amour and Froemke, 2015; Field et al., 2020). In theoretical studies, network models usually include inhibitory plasticity to dynamically stabilize recurrent network dynamics (Vogels et al., 2011; Litwin-Kumar and Doiron, 2014; Zenke et al., 2015). In line with recent efforts to uncover additional functional roles of inhibitory plasticity beyond the stabilization of firing rates (Hennequin et al., 2017), here, we investigated potential functional consequences of inhibitory plasticity in adaptive phenomena. We were inspired by recent experimental work in the mammalian cortex (Chen et al., 2015; Kato et al., 2015; Natan et al., 2015; Hamm and Yuste, 2016; Natan et al., 2017; Heintz et al., 2020), and simpler systems, such as *Aplysia* (Fischer et al., 1997; Ramaswami, 2014) and in *Drosophila* (Das et al., 2011; Glanzman, 2011) along with theoretical reflections (Ramaswami, 2014; Barron et al., 2017), which all point towards a prominent role of inhibition and inhibitory plasticity in the generation of the MMN, SSA, and habituation. For example, Natan and colleagues observed that in the mouse auditory cortex, both parvalbumin-positive (PV) and somatostatin-positive (SOM) interneurons contribute to SSA (Natan et al., 2015), possibly due to inhibitory potentiation (Natan et al., 2017). In the context of habituation, daily passive sound exposure has been found to lead to an up-regulation of the activity of inhibitory neurons (Kato et al., 2015). Furthermore, increased activity to a deviant stimulus in the MMN is diminished when inhibitory neurons are suppressed (Hamm and Yuste, 2016).

Most experimental studies on inhibition in adaptive phenomena have not directly implicated inhibitory plasticity as the relevant mechanism. Instead, some studies have suggested that the firing rate of the inhibitory neurons changes, resulting in more inhibitory input onto excitatory cells, effectively leading to adaptation (Kato et al., 2015). In principle, there can be many other reasons why the inhibitory input increases: disinhibitory circuits, modulatory signals driving specific inhibition, or increased synaptic strength of excitatory to inhibitory cells, to name a few. However, following experimental evidence (Natan et al., 2017) and supported by our results, the plasticity of inhibitory to excitatory connections emerges as a top candidate underlying adaptive phenomena.

One line of evidence to speak against inhibitory plasticity argues that SSA might be independent of NMDA activation (Farley et al., 2010). Inhibitory plasticity, on the contrary, seems to be NMDA receptor-dependent (D’amour and Froemke, 2015; Field et al., 2020). However, there exists some discrepancy in how exactly NMDA receptors are involved in SSA (Ross and Hamm, 2020), since blocking NMDA receptors can disrupt the MMN (Tikhonravov et al., 2008; Chen et al., 2015). These results indicate that a further careful disentanglement of the underlying cellular mechanisms of adaptive phenomena is needed.

### Alternative mechanisms suggested to account for adapted and novelty responses

Undoubtedly, mechanisms other than inhibitory plasticity might underlie the difference in network response to repeated and novel stimuli. These mechanisms can be roughly summarized in two groups: mechanisms which are unspecific, and mechanisms which are specific to the stimulus. Two examples of unspecific mechanisms are intrinsic plasticity and an adaptive current. Intrinsic plasticity is a form of activity-dependent plasticity, adjusting the neurons’ intrinsic excitability (Debanne et al., 2019) and has been suggested to explain certain adaptive phenomena (Levakova et al., 2019). Other models at the single neuron level incorporate an additional current variable, the adaptive current, which increases for each postsynaptic spike and decreases otherwise. This adaptive current leads to a reduction of the neuron’s membrane potential after a spike (Brette and Gerstner, 2005). However, any unspecific mechanism can only account for firing-rate adaptation butnotforSSA(Nelken, 2014)(Fig. 5C). Examples of stimulus-specific mechanisms are short-term plasticity and long-term plasticity of excitatory synapses. Excitatory short-term depression, usually of thalamocortical synapses, is the most widely hypothesized mechanism to underlie adaptive phenomena (Nelken, 2014).

An already established model to explain SSA is the ‘Adaptation of Narrowly Tuned Modules’ (ANTM) model (Nelken, 2014; Khouri and Nelken, 2015), which is largely based on short-term plasticity (Abbott et al., 1997; Tsodyks et al., 1998). This model has been extensively studied in the context of adaptation to tone frequencies (Mill et al., 2011a,b; Taaseh et al., 2011; Mill et al., 2012; Hershenhoren et al., 2014). Models based on short-term plasticity have also been extended to recurrent networks (Yarden and Nelken, 2017) and multiple inhibitory sub-populations (Park and Geffen, 2020). The crucial parameter in adaptation studies based only on short-term depression (as the ANTM model) is the timescale of the short-term dynamics. Experimental studies on adaptation find timescales from hundreds of milliseconds to tens of seconds (Ulanovsky et al., 2004; Lundstrom et al., 2010; Homann et al., 2017; Latimer et al., 2019), and in the case of habituation even multiple days (Haak et al., 2014; Ramaswami, 2014). Although modulation of synaptic strength related to short-term plasticity has been found on the timescales of milliseconds to minutes (Zucker and Regehr, 2002), explaining the different timescales of adaptive phenomena would likely require a dynamic short-term depression timescale. Our work shows that inhibitory plasticity can readily lead to adaptation on multiple timescales without the need for any additional assumptions (Fig. 4). Future work needs to dissect how combining the above-mentioned mechanisms affects adapted and novelty responses.

### Disinhibition as a mechanism for novelty response amplification

Upon including a top-down disinhibitory signal in our model network, we observed: (1) an active amplification of the novelty response (Fig. 6B); (2) elevated responses of unstimulated neurons (Fig. 6C), also referred to as a dense representation (Homann et al., 2017); and (3) a flexible manipulation of neuronal responses through a change in the disinhibitory strength (Fig. 6D, E).

In our model, we were agnostic to the mechanism that generates disinhibition. However, at least two possibilities exist in which the inhibitory population could be regulated by higher-order feedback to allow for disinhibition. First, inhibitory neurons in primary sensory areas can be shaped by diverse neuromodulatory signals, which allow for subtype-specific targeting of inhibitory neurons (Froemke, 2015). Second, higher-order feedback onto layer 1 inhibitory cells could mediate the behavioral relevance of the adapted stimuli through a disinhibitory pathway (Letzkus et al., 2011; Wang and Yang, 2018). Hence, experiments that induce disinhibition either via local mechanisms within the same cortical layer or through higher cortical feedback can provide a test for our postulated role for disinhibition.

In our model, the disinhibitory signal was activated instantaneously. If such additional feedback signals do indeed exist in the brain that signal the detection of higher-order sequence violations, we expect them to arise with a certain delay. Carefully exploring if the dense responses arise with a temporal delay accounting for higher-order processing and projection back to primary sensory areas might shed light on distributed computations upon novel stimuli. These experiments would probably require recording methods on a finer temporal scale than calcium imaging.

Experimental data which points towards a flexible modulation of novelty and adapted responses already exists. The active amplification of novelty responses generated by our model is consistent with some experimental data (Taaseh et al., 2011; Hershenhoren et al., 2014; Hamm and Yuste, 2016; Harms et al., 2016), but see also (Vinken et al., 2017). Giving a behavioral meaning to a sound through fear conditioning has been shown to modify SSA (Yaron et al., 2020). Similarly, contrast adaptation has been shown to reverse when visual stimuli become behaviorally relevant(Keller et al., 2017). Other studies have also shown that as soon as a stimulus becomes behaviorally relevant, inhibitory neurons decrease their response and therefore disinhibit adapted excitatory neurons (Kato et al., 2015; Makino and Komiyama, 2015; Hattori et al., 2017). Attention might lead to activation of the disinhibitory pathway, allowing for a change in the novelty response compared to the unattended case, as suggested in MMN studies (Sussman et al., 2014). Especially in habituation, the idea that a change in context can assign significance to a stimulus and therefore block habituation, leading to ‘dehabituation’, is widely accepted (Ramaswami, 2014; Barron et al., 2017).

Hence we suggest that disinhibition is a flexible mechanism to control several aspects of novelty responses, including the density of the response, which might be computationally important in signaling change detection to downstream areas (Homann et al., 2017). Altogether, our results suggest that disinhibition is capable of accounting for various aspects of novelty responses that cannot be accounted for by bottom-up computations. The functional purpose of a dense response to novelty stimuli are yet to be explored.

### Functional implications of adapted and novelty responses

In theoretical terms, our model is an attractor network. It differs from classic attractor models where inhibition is considered unspecific (like a ‘blanket’) (Amit and Brunel, 1997). Computational work is starting to uncover the functional role of specific inhibition in static networks (Rost et al., 2018; Najafi et al., 2020; Rostami et al., 2020) as well as the plasticity mechanisms that allow for specific connectivity to emerge (Mackwood et al., 2020). These studies have argued that inhibitory assemblies can improve the robustness of attractor dynamics (Rost et al., 2018) and keep a local balance of excitation and inhibition (Rostami et al., 2020). We showed that specific inhibitory connections readily follow from a tuned inhibitory population (Fig. 1A, Suppl. Fig. S1). Our results suggest that adaptation is linked to a stimulus-specific excitatory/inhibitory (E/I) balance. Presenting a novel stimulus leads to a short-term disruption of the E/I balance, triggering inhibitory plasticity, which aims to restore the E/I balance (Fig. 4) (Vogels et al., 2011; D’amour and Froemke, 2015; Field et al., 2020). Disinhibition, which effectively disrupts the E/I balance, allows for flexible control of adapted and novelty responses (Fig. 6). This links to the notion of disinhibition as a gating mechanism for learning and plasticity (Froemke et al., 2007; Letzkus et al., 2011; Kuhlman et al., 2013).

A multitude of functional implications have been suggested for the role of adaptation (Weber et al., 2019; Snow et al., 2017). We showed that one of these roles, the detection of unexpected (or novel) events, follows from the lack of selective adaptation to those events. A second, highly considered functional implication is predictive coding. In the predictive coding framework, the brain is viewed as an inference or a prediction machine. It is thought to generate internal models of the world which are compared to the incoming sensory inputs (Bastos et al., 2012; Clark, 2013; Friston, 2018). According to predictive coding, the overall goal of our brain is to minimize the prediction error, i.e. the difference between the internal prediction and the sensory input (Rao and Ballard, 1999; Clark, 2013; Friston, 2018). Most predictive coding schemes hypothesize the existence of two populations of neurons. First, prediction error units that signal a mismatch between the internal model prediction and the incoming sensory stimuli. And second, a prediction population unit that reflects what the respective layer ‘knows about the world’ (Rao and Ballard, 1999; Clark, 2013; Spratling, 2017). Our model suggests that primary sensory areas allow for bottom-up detection of stimulus changes without the need for an explicit population of error neurons or an internal model of the world. However, one could also interpret the state of all inhibitory synaptic weights as an implicit internal model of the recent frequency of various events in the environment.

### Predictions and Outlook

Our approach to mechanistically understand the generation of adapted and novelty responses leads to several testable predictions. First, the most general implication from our study is that inhibitory plasticity might serve as an essential mechanism underlying many adaptive phenomena. Our work suggests that inhibitory plasticity allows for adaptation on multiple timescales, ranging from the adaptation to sequence blocks on the timescale of seconds to slower adaptation on the timescale of minutes, corresponding to repeating multiple sequence blocks (Fig. 4C, D). A second prediction follows from the finding that both excitatory and inhibitory neuron populations show adaptive behavior and novelty responses (Fig. 3B, C). Adaptation of inhibitory neurons on the single-cell level has already been verified experimentally (Chen et al., 2015; Natan et al., 2015). Third, we further predict that a violation of the sequence order does not lead to a novelty response. Therefore, the novelty response should not be interpreted as signaling a violation of the exact temporal structure of the sequence (Fig. 3D, E). Furthermore, the height of the novelty peak in the population average depends on the input drive, where decreasing the input strength decreases the novelty response (Fig. 3F). This could be tested, e.g., in the visual system, by presenting visual stimuli with different contrasts.

In our modeling approach, we did not distinguish between different subtypes of inhibitory neurons. This assumption is certainly an oversimplification. The main types of inhibitory neurons, somatostatin-positive (SOM), parvalbumin-positive (PV), and vasoactive intestinal peptide (VIP) expressing neurons, differ in their connectivity and their hypothesized functional roles (Tremblay et al., 2016). This is certainly also true for adaptation, and computational studies have already started to tackle this problem (Park and Geffen, 2020). Studies of the influence of inhibitory neurons on adaptation have shown that different interneuron types have unique contributions to adaptation (Kato et al., 2015; Natan et al., 2015; Hamm and Yuste, 2016; Natan et al., 2017; Garrett et al., 2020; Heintz et al., 2020). It would be interesting to combine the knowledge of microcircuit connectivity of excitatory neurons, PVs, SOMs, and VIPs with subtype-specific inhibitory plasticity mechanisms (Agnes et al., 2019).

In sum, we have proposed a mechanistic model for the emergence of adapted and novelty responses based on inhibitory plasticity, and the regulation of this novelty response by top-down signals. Our findings offer insight into the flexible and adaptive responses of animals in constantly changing environments, and could be further relevant for disorders like schizophrenia where adaptive responses are perturbed (Hamm et al., 2017).

## Methods

We built a biologically-plausible spiking neuronal network model of the mammalian cortex based on recent experimental findings on tuning, connectivity, and synaptic plasticity. The model consisted of 4000 excitatory exponential integrate-and-fire (EIF) neurons and 1000 inhibitory leaky integrate-and-fire (LIF) (Table 1). Excitatory (E) and inhibitory (I) neurons were randomly recurrently connected (Table 2). Plasticity from excitatory-to-excitatory neurons was based on excitatory triplet spike-timing-dependent plasticity (eSTDP) rule, which uses triplets of pre- and post-synaptic spikes to evoke synaptic change and captures the firing rate dependency found experimentally (Sjöström et al., 2001; Pfister and Gerstner, 2006; Gjorgjieva et al., 2011) (Table3). In addition, excitatory-to-excitatory weight dynamics were stabilized by a heterosynaptic plasticity mechanism (Fiete et al., 2010; Field et al., 2020), which preserved the total sum of all incoming synaptic weights into an excitatory neuron. Plasticity from inhibitory to excitatory neurons was modeled based on an inhibitory pairwise STDP (iSTDP) rule, which allows stabilization of excitatory firing rate dynamics (Vogels et al., 2011; Litwin-Kumar and Doiron, 2014) and has been confirmed experimentally (D’amour and Froemke, 2015). All other synapses in the network are fixed. Both excitatory and inhibitory neurons received an excitatory baseline feedforward input in the form of Poisson spikes. Furthermore, different subsets of excitatory and inhibitory neurons received excess input with elevated Poisson rate (Fig. 1A, left; Table 4). Inhibitory tuning in the model was assumed to be broader than excitatory tuning, reflected in the larger probability of an inhibitory neuron to be driven by a stimulus than an excitatory neuron (Table 4). Stimulus tuning in both populations led to the formation of stimulus-specific excitatory assemblies, where the subsets of excitatory neurons receiving the same input developed strong connections among each other (Suppl. Fig. S1), as found experimentally (Ko et al., 2011, 2013; Miller et al., 2014; Lee et al., 2016). Additionally, the connections from similarly tuned inhibitory to excitatory neurons also became stronger, as seen in experiments (Lee et al., 2014; Xue et al., 2014; Znamenskiy et al., 2018; Najafi et al., 2020). The number of stimulus-specific assemblies varies depending on the stimulation paradigm and corresponds to the number of unique stimuli presented in a given paradigm. One neuron can be driven by multiple stimuli. This means that stimulus-specific assemblies can have significant overlap.

**Table 1.** Parameters for the excitatory (EIF) and inhibitory (LIF) membrane dynamics (Litwin-Kumar and Doiron, 2014).

| Symbol | Description | Value |
| --- | --- | --- |
| $N^E$ | Number of E neurons | 4,000 |
| $N^I$ | Number of I neurons | 1,000 |
| $\tau^E, \tau^I$ | E, I neuron resting membrane time constant | 20 ms |
| $V_{rest}^E$ | E neuron resting potential | - 70 mV |
| $V_{rest}^I$ | I neuron resting potential | - 62 mV |
| $\Delta_T$ | Slope factor of exponential | 2 mV |
| $C$ | Membrane capacitance | 300 pF |
| $g_L$ | Membrane conductance | $C/\tau^E$ |
| $V_{rev}^E$ | E reversal potential | 0 mV |
| $V_{rev}^I$ | I reversal potential | - 75 mV |
| $V_{thr}$ | Threshold potential | - 52 mV |
| $V_{peak}$ | Peak threshold potential | 20 mV |
| $V_{reset}$ | E, I neuron reset potential | - 60 mV |
| $\tau_{abs}$ | E, I absolute refractory period | 1 ms |

### Dynamics of synaptic conductances and the membrane potential

The membrane dynamics of each excitatory neuron was modeled as an exponential integrate-and-fire (EIF) neuron model (Fourcaud-Trocmé et al., 2003):

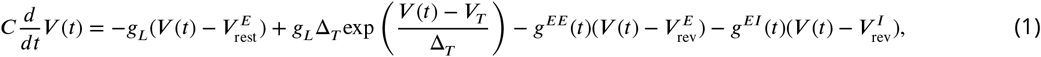

where *V*(*t*) is the membrane potential of the modeled neuron, *C* the membrane capacitance, *g_L_* the membrane conductance, and Δ_*T*_ is the slope factor of the exponential rise. The membrane potential was reset to *V*_reset_ once the diverging potential reached the threshold peak voltage *V*_peak_. Inhibitory neurons were modeled via a leaky-integrate-and-fire neuron model

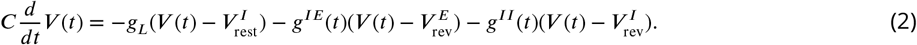

Once the membrane potential reached the threshold voltage *V*_thr_, the membrane potential was reset to *V*_reset_. The absolute refractory period was modeled by clamping the membrane voltage of a neuron that just spiked to the reset voltage *V*_reset_ for the duration *τ*_abs_. In this manuscript we did not model additional forms of adaptation, such as adaptive currents or spiking threshold *V*_T_ adaptation. To avoid extensive parameter tuning, we used previously published parameter values (Litwin-Kumar and Doiron, 2014)(Table 1).

We compare this model to one where we freeze plasticity and include adaptive currents *w*_adapt_ (Fig. 5B, top). We model this by subtracting *w*_adapt_(*t*) on the right hand side of Eq. 1 (Brette and Gerstner, 2005). Upon a spike, *w*_adapt_(*t*) increases by *b_w_* and the sub-threshold dynamics of the adaptive current are described by 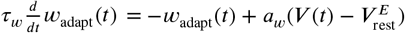, where *a_w_* = 4 nS denotes the subthreshold and *b_w_* = 80.5 pA the spike-triggered adaptation. The adaptation time scale is set to *τ_w_* = 150 ms.

The conductance of neuron *i* which is part of population *X* and is targeted by another neuron in population *Y* is denoted with 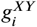. Both *X* and *Y* can refer either to the excitatory or inhibitory population, i.e. *X, Y* ∈ [*E, I*]. The shape of the synaptic kernels *F*(*t*) is a difference of exponentials and differs for excitatory and inhibitory input depending on the rise and decay times 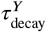 and 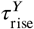:

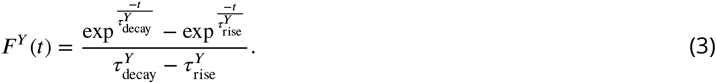

This kernel is convolved with the total inputs to neuron *i* weighted with the respective synaptic strength to yield the total conductance

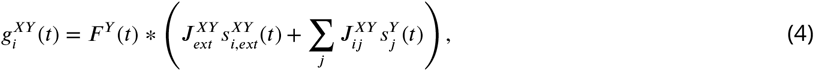

where 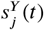 is the spike train of neuron *j* in the network with index and 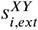 denotes the spike train of the external input to neuron *i*. The external spike trains are generated in an independent homogeneous Poisson process. The synaptic strength from the input units to the network neurons, 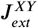, is assumed to be constant.

### Excitatory and inhibitory plasticity

We chose to implement the plasticity from an excitatory to an excitatory neuron *J^EE^* based on the triplet spike time dependent plasticity rule (triplet STDP) (Pfister and Gerstner, 2006). In the triplet rule, four spike accumulators, *r*_1_, *r*_2_, *o*_1_, and *o*_2_, increase by one, once a spike of the corresponding neuron occurs and otherwise decrease exponentially depending on their respective time constant *τ*_+_, *τ_x_*, *τ*_−_, and *τ_y_*:

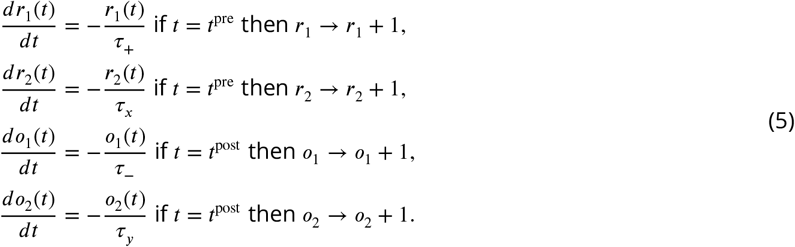

The E to E weights are updated as

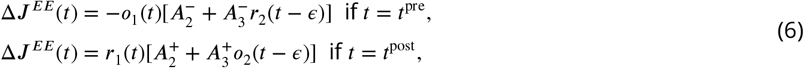

where the *A*^+^, *A*^−^ corresponds to the excitatory LTP or LTD amplitude, and the subscript refers to the triplet (3) or pairwise term (2). The parameter *ϵ* > 0 ensures that the weights are updated prior to increasing the respective spike accumulators by 1. Spike detection is modeled in an all-to-all approach.

For inhibitory to excitatory neuron connections, *J^EI^* the inhibitory spike-timing-dependent plasticity rule (iSTDP) is implemented (Vogels et al., 2011). The plasticity parameters are shown in table 3. The two spike accumulators *y^E/I^*, for the inhibitory pre and the excitatory post-synaptic neuron, have the same time constant 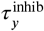. Their dynamics are described by

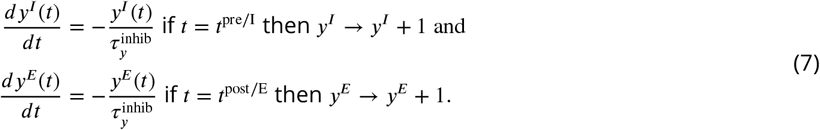

The I to E weights are updated as

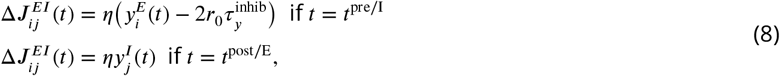

where *η* is the learning rate, and *r*_0_ corresponds to the target firing rate of the excitatory neuron.

### Additional homeostatic mechanisms in recurrent networks

Inhibitory plasticity alone is considered insufficient to prevent runaway activity in this network implementation. Hence, additional mechanisms were implemented that also have a homeostatic effect. To further avoid unbound increasing weights, the synaptic weights are only allowed to change within bounds, the exact parameters can be found in table 2. Subtractive normalization ensures that the total synaptic input to an excitatory neuron remains constant throughout the simulation. This is implemented by scaling all incoming weights to each neuron every 20 s according to

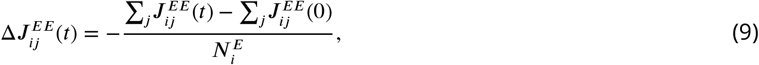

where *i* is the index of the post-synaptic and *j* of the pre-synaptic neurons. 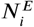 is the number of excitatory connections onto neuron *i*.

**Table 2.** Parameters for feedforward and recurrent connections (Litwin-Kumar and Doiron, 2014).

| Symbol | Description | Value |
| --- | --- | --- |
| $p$ | Connection probability | 0.2 |
| $\tau_{\text{rise}}^E$ | Rise time for E synapses | 1 ms |
| $\tau_{\text{decay}}^E$ | Decay time for E synapses | 6 ms |
| $\tau_{\text{rise}}^I$ | Rise time for I synapses | 0.5 ms |
| $\tau_{\text{decay}}^I$ | Decay time for I synapses | 2 ms |
| $\bar{r}_{\text{ext}}^{EE}$ | Avg. rate of external input to E neurons | 4.5 kHz |
| $\bar{r}_{\text{ext}}^{IE}$ | Avg. rate of external input to I neurons | 2.25 kHz |
| $J_{\text{min}}^{EE}$ | Minimum E to E synaptic weight | 1.78 pF |
| $J_{\text{max}}^{EE}$ | Maximum E to E synaptic weight | 21.4 pF |
| $J_0^{EE}$ | Initial E to E synaptic weight | 2.76 pF |
| $J_{\text{min}}^{EI}$ | Minimum I to E synaptic weight | 48.7 pF |
| $J_{\text{max}}^{EI}$ | Maximum I to E synaptic weight | 243 pF |
| $J_0^{EI}$ | Initial I to E synaptic weight | 48.7 pF |
| $J^{IE}$ | Synaptic weight from E to I | 1.27 pF |
| $J^{II}$ | Synaptic weight from I to I | 16.2 pF |
| $J^{EE\text{x}}$ | Synaptic weight from external input population to E | 1.78 pF |
| $J^{IE\text{x}}$ | Synaptic weight from external input population to I | 1.27 pF |

**Table 3.** Parameters for the implementation of Hebbian and homeostatic plasticity (Pfister and Gerstner, 2006; Litwin-Kumar and Doiron, 2014).

| Symbol | Description | Value |
| --- | --- | --- |
| $\tau_-$ | Time constant of pairwise pre-synaptic detector (+) | 33.7 ms |
| $\tau_+$ | Time constant of pairwise post-synaptic detector (-) | 16.8 ms |
| $\tau_x$ | Time constant of triplet pre-synaptic detector (-) | 101 ms |
| $\tau_y$ | Time constant of triplet post-synaptic detector (+) | 125 ms |
| $A_2^+$ | Pairwise potentiation amplitude | $7.5 \times 10^{-10}$ pF |
| $A_3^+$ | Triplet potentiation amplitude | $9.3 \times 10^{-3}$ pF |
| $A_2^-$ | Pairwise depression amplitude | $7 \times 10^{-3}$ pF |
| $A_3^-$ | Triplet depression amplitude | $2.3 \times 10^{-4}$ pF |
| $\tau_y^{\text{inhib}}$ | Time constant of low-pass filtered spike train | 20 ms |
| $\eta$ | Inhibitory plasticity learning rate | 1 pF |
| $r_0$ | Target firing rate | 3 Hz |
| $\Delta t_{\text{het}}$ | Heterosynaptic plasticity normalisation | 20 ms |

### Pretraining phase

During the pretraining phase, i.e. before the stimulation paradigm starts, we sequentially stimulate the groups of neurons that are driven by the stimuli (including’novel’ stimuli) in the consecutive stimulation protocol. The order is random, leading to a change in network connectivity that is only stimulus but not sequence-dependent (Fig. 4B, first 100 s shown here for five repetitions of each stimulus). The pretraining phase is a phenomenological model of the development process to reach some structure in the connections prior to the actual stimulation paradigm.

### Stimulation protocol

All neurons receive external excitatory baseline input. The baseline input to excitatory neurons 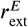 is higher than the input to inhibitory neurons 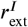 (Table 4). An external input of 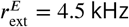 relates to 1000 external pre-synaptic neurons with average firing rates of 4.5 Hz (compare Litwin-Kumar and Doiron (2014)). The model analog of the presentation of a sensory stimulus in experiments is increased input to a subset of neurons. Every time a particular stimulus is repeated, the same set of neurons receives stronger external stimulation, 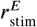 and 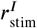. In contrast to several previous plastic recurrent networks, we do not only consider the excitatory neurons to have stimulus tuning properties but include inhibitory tuning as well. The probability of an excitatory neuron to be driven by one particular stimulus is 5%, leading to roughly 200 neurons that are specific to this stimulus. We model inhibitory tuning to be both weaker and more broad. The probability of an inhibitory neuron to be driven by one particular stimulus is 15%, leading to roughly 150 neurons that respond specifically to this stimulus. There is overlap in stimulus tuning, i.e. one neuron can be driven by multiple stimuli. The stimulation paradigm was inspired by a recent study in the visual system (Homann et al., 2017). The timescales of the experimental paradigms and the model are matched, i.e. the neurons tuned to a stimulus receive additional input for 300 ms simulation time. Stimuli are presented without pauses in between, corresponding to continuous stimulus presentation without blank images (visual) or silence (auditory) between sequence blocks. The stimulus parameters are listed in table 4. Disinhibition in the model is implemented via additional inhibiting input to the inhibitory population 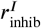. This is modeled purely in a phenomenological way, and we are agnostic as to what causes the additional inhibition.

**Table 4.** Parameters for the stimulation paradigm and stimulus tuning.

| Symbol | Description | Value |
| --- | --- | --- |
| $r_{\text{ext}}^E$ | External baseline input to E | 4.5 kHz |
| $r_{\text{ext}}^I$ | External baseline input to I | 2.25 kHz |
| $r_{\text{stim}}^E$ | Additional input to E during stimulus presentation | 12 kHz |
| $r_{\text{stim}}^I$ | Additional input to I during stimulus presentation | 1.2 kHz |
| $r_{\text{disinh}}^I$ | Additional input to I during disinhibition | -1.5 kHz |
| $p_{\text{member}}^E$ | Probability for an E neuron to be driven by a stimulus | 5% |
| $p_{\text{member}}^I$ | Probability for an I neuron to be driven by a stimulus | 15% |

The simulations were performed using the Julia programming language. Further evaluation and plotting was done in Python. Euler integration was implemented using a time step of 0.1 ms. Code implementing our model and generating the stimulation protocols will be made available upon publication.

## Acknowledgements

AS, CM and JG thank the Max Planck Society for funding and MJB thanks the NEI and the Princeton Accelerator Fund for funding. We thank members of the ‘Computation in Neural Circuits’ group for useful discussions and comments on the manuscript.

---

**Figure S1.**
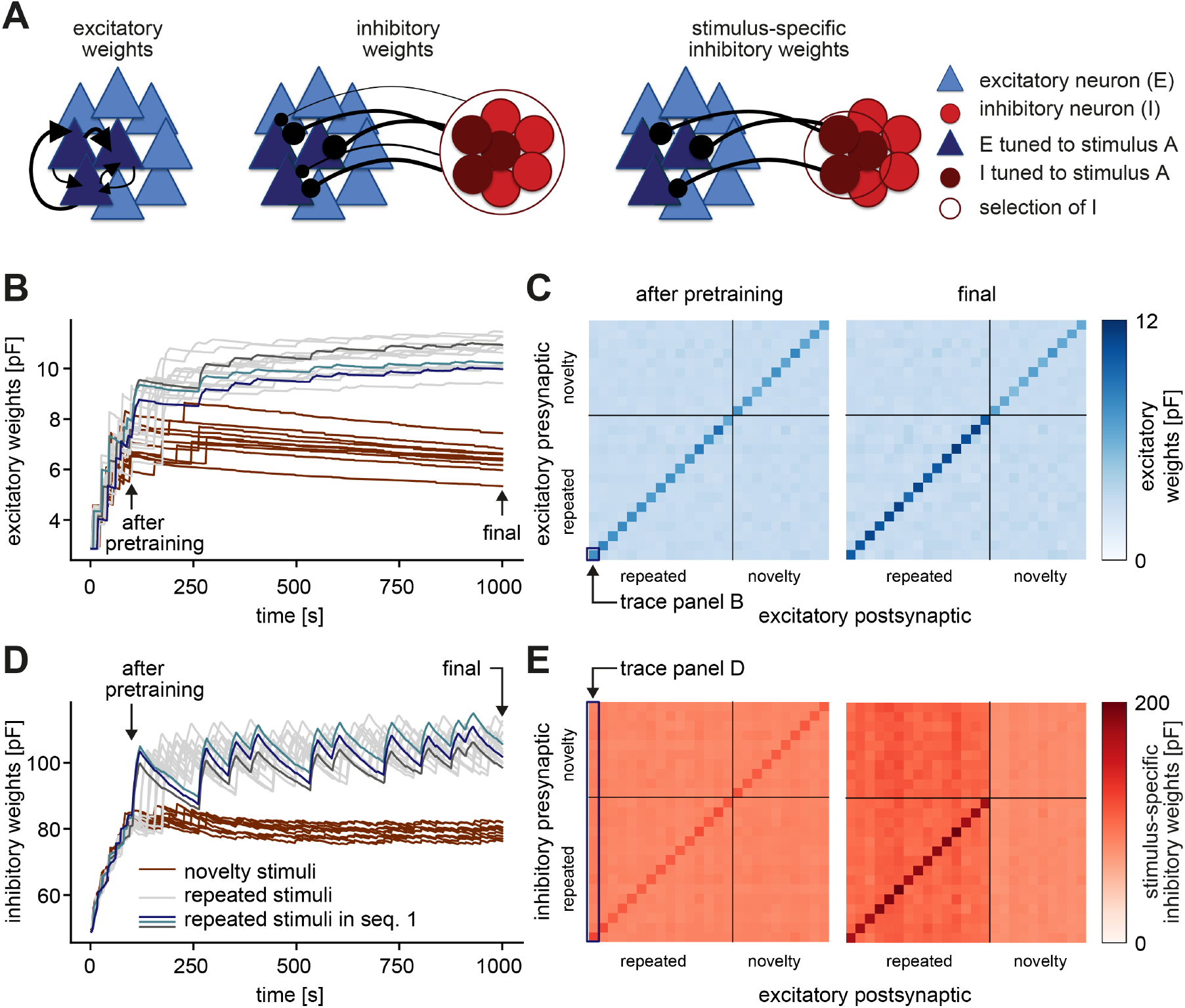
Strong connections form between excitatory and excitatory, as well as inhibitory and excitatory neuron groups that are tuned to the same stimulus. **A.** A recurrently connected network of excitatory (E) neurons (blue triangles) and inhibitory (I) neurons (red circles) receives tuned input. Excitatory neurons tuned to a sample stimulus A are highlighted in dark blue, the inhibitory counterparts in dark red. Left: Average excitatory weights within stimulus-specific assembly A are determined by averaging all E-to-E weights within an assembly. Center: Average inhibitory weights onto stimulus-specific assembly A are determined by averaging the weights from all inhibitory neurons onto stimulus-specific assembly A. Right: Stimulus-specific inhibitory weights onto stimulus-specific assembly A are determined by averaging only the weights from the inhibitory neurons that are also tuned to stimulus A (dark red). **B.** Evolution of the average excitatory weights corresponding to all repeated and 10 sample novelty stimuli. Colored traces mark three stimulus-specific assemblies in sequence 1: A, B, and C. The legend is shared with panel D. **C.** Left: Weight matrix of average excitatory (E-to-E) weights after pretraining (see panel B) for all repeated and 10 sample novelty stimuli separated by the gray lines. The size of assemblies can be slightly different. The weights across assemblies (off-diagonal) are small compared to the weights within assemblies (diagonal). After pretraining, there is no apparent difference in connectivity for novel and repeated stimuli. The diagonal elements correspond to the traces plotted in B, i.e. the time evolution of the bottom-left square, highlighted in dark blue corresponds to the dark blue trace in panel B. Right: Same as left after the repeated sequence stimulation paradigm (see final in panel B). Here, assembly weights for repeated stimuli are stronger. **D.** Same as panel B for the average inhibitory weights. Arrows indicate end points of the pretraining and whole stimulation phase. **E.** Left: Weight matrix of stimulus-specific inhibitory (I-to-E) weights after pretraining (see panel D) for all repeated and 10 sample novelty stimuli separated by the gray lines. The size of assemblies can be slightly different. Inhibition is stimulus-specific (diagonal stronger than off diagonal), i.e. the weights from inhibitory neurons tuned to a given stimulus onto excitatory neurons that are tuned to the same stimulus (diagonal) are larger that onto excitatory neurons that are tuned to different stimuli (off-diagonal). After pretraining, there is no apparent difference in connectivity for novel and repeated stimuli. Here, an entire column average approximately corresponds to the traces plotted in D, i.e. the time evolution of the averaged first column, highlighted in dark blue corresponds to the dark blue trace in panel D. It is only approximate, since individual inhibitory neurons can be tuned to multiple repeated and novel stimuli and hence contribute to multiple averages shown here in the matrix. Right: Same as left after the repeated sequence stimulation paradigm (see final in panel D). Here, excitatory assemblies tuned to repeated stimuli receive more inhibition than those tuned to novel stimuli.

## References

Abbott LF, Varela JA, Sen K, Nelson SB. Synaptic Depression and Cortical Gain Control. Science. 1997; 275(5297):221–224. doi: https://doi.org/10.1126/science.275.5297.221.

Agnes EJ, Luppi AI, Vogels TP. Complementary inhibitory receptive fields emerge from synaptic plasticity and create an attentional switch in sensory circuits. bioRxiv. 2019; 729988. doi: https://doi.org/10.1101/729988.

Amit DJ, Brunel N. Model of Global Spontaneous Activity and Local Structured Activity During Delay Periods in the Cerebral Cortex. Cerebral Cortex. 1997; 7:237–252. doi: https://doi.org/10.14257/ijbsbt.2015.7.5.05.

Barlow HB. Possible Principles Underlying the Transformations of Sensory Messages. Sensory Communication. 2013; 1:216–234. doi: https://doi.org/10.7551/mitpress/9780262518420.003.0013.

Barron HC, Vogels TP, Behrens TE, Ramaswami M. Inhibitory engrams in perception and memory. Proceedings of the National Academy of Sciences. 2017; 114(26):6666–6674. doi: https://doi.org/10.1073/pnas.1701812114.

Bastos AM, Usrey WM, Adams RA, Mangun GR, Fries P, Friston KJ. Canonical Microcircuits for Predictive Coding. Neuron. 2012; 76(4):695–711. doi: https://doi.org/10.1016/j.neuron.2012.10.038.

Brette R, Gerstner W. Adaptive Exponential Integrate-and-Fire Model as an Effective Description of Neuronal Activity. Journal of Neurophysiology. 2005; 94:3637–3642. doi: https://doi.orcheng/10.1152/jn.00686.2005.

Chen IW, Helmchen F, Lütcke H. Specific early and late oddball-evoked responses in excitatory and inhibitory neurons of mouse auditory cortex. Journal of Neuroscience. 2015; 35(36):12560–12573. doi: https://doi.org/10.1523/JNEUROSCI.2240-15.2015.

Clark A. Whatever next? Predictive brains, situated agents, and the future of cognitive science. Behavioral and Brain Sciences. 2013; 36:181–253. doi: https://doi.org/10.1017/S0140525X12000477.

D’amour JA, Froemke RC. Inhibitory and Excitatory Spike-Timing-Dependent Plasticity in the Auditory Cortex. Neuron. 2015; 86:514–528. doi: https://doi.org/10.1016/j.neuron.2015.03.014.

Das S, Sadanandappa MK, Dervan A, Larkin A, Lee JA, Sudhakaran IP, Priya R, Heidari R, Holohan EE, Pimentel A, Gandhi A, Ito K, Sanyal S, Wang JW, Rodrigues V, Ramaswami M. Plasticity of local GABAergic interneurons drives olfactory habituation. Proceedings of the National Academy of Sciences of the United States of America. 2011; 108(36):E646–E654. doi: https://doi.org/10.1073/pnas.1106411108.

Debanne D, Inglebert Y, Russier M. Plasticity of intrinsic neuronal excitability. Current Opinion in Neurobiology. 2019; 54:73–82. doi: https://doi.org/10.1016/j.conb.2018.09.001.

Dhruv NT, Carandini M. Cascaded Effects of Spatial Adaptation in the Early Visual System. Neuron. 2014; 81:529–535. doi: https://doi.org/10.1016/j.neuron.2013.11.025.

Fairhall AL. Adaptation and natural stimulus statistics. In: Gazzaniga MS, Mangun GR, editors. The Cognitive Neurosciences, 5 ed. MIT Press; 2014. p. 283–294.

Farley BJ, Quirk MC, Doherty JJ, Christian EP. Stimulus-specific adaptation in auditory cortex is an NMDA-independent process distinct from the sensory novelty encoded by the mismatch negativity. Journal of Neuroscience. 2010; 30(49):16475–16484. doi: https://doi.org/10.1523/JNEUROSCI.2793-10.2010.

Field RE, D’amour JA, Tremblay R, Miehl C, Rudy B, Gjorgjieva J, Froemke RC. Heterosynaptic Plasticity Determines the Set Point for Cortical Excitatory-Inhibitory Balance. Neuron. 2020; 106(5):842–854. doi: https://doi.org/10.1016/j.neuron.2020.03.002.

Fiete IR, Senn W, Wang CZH, Hahnloser RHR. Spike-Time-Dependent Plasticity and Heterosynaptic Competition Organize Networks to Produce Long Scale-Free Sequences of Neural Activity. Neuron. 2010; 65:563–576. doi: https://doi.org/10.1016/j.neuron.2010.02.003.

Fischer TM, Blazis DEJ, Priver NA, Carew TJ. Metaplasticity at identified inhibitory synapses in Aplysia. Nature. 1997; 389:860–865. doi: https://doi.org/10.1038/39892.

Fourcaud-Trocmé N, Hansel D, van Vreeswijk C, Brunel N. How spike generation mechanisms determine the neuronal response to fluctuating inputs. The Journal of Neuroscience. 2003; 23(37):11628–11640. doi: https://doi.org/10.1523/JNEUROSCI.23-37-11628.2003.

Friston K. Does predictive coding have a future? Nature Neuroscience. 2018; 21:1019–1021. doi: https://doi.org/10.1038/s41593-018-0200-7.

Froemke RC. Plasticity of Cortical Excitatory-Inhibitory Balance. Annual Review of Neuroscience. 2015; 38:195–219. doi: https://doi.org/10.1146/annurev-neuro-071714-034002.

Froemke RC, Merzenich MM, Schreiner CE. A synaptic memory trace for cortical receptive field plasticity. Nature. 2007; 450:425429. doi: https://doi.org/10.1038/nature06289.

Garrett M, Manavi S, Roll K, Ollerenshaw DR, Groblewski PA, Ponvert ND, Kiggins JT, Casal L, Mace K, Williford A, Leon A, Jia X, Ledochowitsch P, Buice MA, Wakeman W, Mihalas S, Olsen SR. Experience shapes activity dynamics and stimulus coding of VIP inhibitory cells. eLife. 2020; 9:e50340. doi: https://doi.org/10.7554/eLife.50340.

Geffen MN, deVries SEJ, Meister M. Retinal ganglion cells can rapidly change polarity from off to on. PLoS Biology. 2007; 5(3):e65. doi: https://doi.org/10.1371/journal.pbio.0050065.

Gjorgjieva J, Clopath C, Audet J, Pfister JP. A triplet spike-timing-dependent plasticity model generalizes the Bienenstock-Cooper-Munro rule to higher-order spatiotemporal correlations. Proceedings of the National Academy of Sciences. 2011; 108(48):19383–19388. doi: https://doi.org/10.1073/pnas.1105933108.

Glanzman DL. Olfactory habituation: Fresh insights from flies. Proceedings of the National Academy of Sciences of the United States of America. 2011; 108(36):14711–14712. doi: https://doi.org/10.1073/pnas.1111230108.

Griffen TC, Maffei A. GABAergic synapses: their plasticity and role in sensory cortex. Frontiers in Cellular Neuroscience. 2014; 8(91). doi: https://doi.org/10.3389/fncel.2014.00091.

Haak KV, Fast E, Bao M, Lee M, Engel SA. Four days of visual contrast deprivation reveals limits of neuronal adaptation. Current Biology. 2014; 24(21):2575–2579. doi: https://doi.org/10.1016/j.cub.2014.09.027.

Haas JS, Nowotny T, Abarbanel HDI. Spike-Timing-Dependent Plasticity of Inhibitory Synapses in the Entorhinal Cortex. Journal of Neurophysiology. 2006; 96:3305–3313. doi: https://doi.org/10.1152/jn.00551.2006.

Hamm JP, Peterka DS, Gogos JA, Yuste R. Altered Cortical Ensembles in Mouse Models of Schizophrenia. Neuron. 2017; 94:153167. doi: http://dx.doi.org/10.1016/j.neuron.2017.03.019.

Hamm JP, Yuste R. Somatostatin Interneurons Control a Key Component of Mismatch Negativity in Mouse Visual Cortex. Cell Reports. 2016; 16(3):597–604. doi: https://doi.org/10.1016/j.celrep.2016.06.037.

Harms L, Michie PT, Näätänen R. Criteria for determining whether mismatch responses exist in animal models: Focus on rodents. Biological Psychology. 2016; 116:28–35. doi: http://dx.doi.org/10.1016/j.biopsycho.2015.07.006.

Hattori R, Kuchibhotla KV, Froemke RC, Komiyama T. Functions and dysfunctions of neocortical inhibitory neuron subtypes. Nature Neuroscience. 2017; 20(9):1199–1208. doi: https://doi.org/10.1038/nn.4619.

Heintz TG, Hinojosa AJ, Lagnado L. Opposing forms of adaptation in mouse visual cortex are controlled by distinct inhibitory microcircuits and gated by locomotion. bioRxiv. 2020; 909788. doi: https://doi.org/10.1101/2020.01.16.909788.

Hennequin G, Agnes EJ, Vogels TP. Inhibitory Plasticity: Balance, Control, and Codependence. Annual Review of Neuroscience. 2017; 40:557–579. doi: https://doi.org/10.1146/annurev-neuro-072116-031005.

Hershenhoren I, Taaseh N, Antunes FM, Nelken I. Intracellular correlates of stimulus-specific adaptation. Journal of Neuroscience. 2014; 34(9):3303–3319. doi: https://doi.org/10.1523/JNEUROSCI.2166-13.2014.

Holmgren CD, Zilberter Y. Coincident Spiking Activity Induces Long-Term Changes in Inhibition of Neocortical Pyramidal Cells. The Journal of Neuroscience. 2001; 21(20):8270–8277. doi: https://doi.org/10.1523/jneurosci.21-20-08270.2001.

Homann J, Koay SA, Glidden AM, Tank DW, Berry II MJ. Predictive Coding of Novel versus Familiar Stimuli in the Primary Visual Cortex. bioRxiv. 2017; 197608. doi: https://doi.org/10.1101/197608.

Kato HK, Gillet SN, Isaacson JS. Flexible Sensory Representations in Auditory Cortex Driven by Behavioral Relevance. Neuron. 2015; 88(5):1027–1039. doi: http://dx.doi.org/10.1016/j.neuron.2015.10.024.

Keller AJ, Houlton R, Kampa BM, Lesica NA, Mrsic-Flogel TD, Keller GB, Helmchen F. Stimulus relevance modulates contrast adaptation in visual cortex. eLife. 2017; 6:e21589. doi: https://doi.org/10.7554/eLife.21589.

Keller GB, Bonhoeffer T, Hübener M. Sensorimotor Mismatch Signals in Primary Visual Cortex of the Behaving Mouse. Neuron. 2012; 74(5):809–815. doi: https://doi.org/10.1016/j.neuron.2012.03.040.

Khouri L, Nelken I. Detecting the unexpected. Current Opinion in Neurobiology. 2015; 35:142–147. doi: http://dx.doi.org/10.1016/j.conb.2015.08.003.

King JL, Lowe MP, Stover KR, Wong AA, Crowder NA. Adaptive processes in Thalamus and cortex revealed by silencing of primary visual cortex during contrast adaptation. Current Biology. 2016; 26(10):1295–1300. doi: http://dx.doi.org/10.1016/j.cub.2016.03.018.

Ko H, Cossell L, Baragli C, Antolik J, Clopath C, Hofer SB, Mrsic-Flogel TD. The emergence of functional microcircuits in visual cortex. Nature. 2013; 496:96–100. doi: https://doi.org/10.1038/nature12015.

Ko H, Hofer SB, Pichler B, Buchanan KA, Sjöström PJ, Mrsic-Flogel TD. Functional specificity of local synaptic connections in neocortical networks. Nature. 2011; 473:87–91. doi: https://doi.org/10.1038/nature09880.

Kuhlman SJ, Olivas ND, Tring E, Ikrar T, Xu X, Trachtenberg JT. A disinhibitory microcircuit initiates critical-period plasticity in the visual cortex. Nature. 2013; 501:543–546. doi: https://doi.org/10.1038/nature12485.

Latimer K, Barbera D, Sokoletsky M, Awwad B, Katz Y, Nelken I, Lampl I, Fairhall AL, Priebe NJ. Multiple timescales account for adaptive responses across sensory cortices. Journal of Neuroscience. 2019; 39(50):10019–10033. doi: https://doi.org/10.1523/JNEUROSCI.1642-19.2019.

Lee SH, Marchionni I, Bezaire M, Varga C, Danielson N, Lovett-Barron M, Losonczy A, Soltesz I. Parvalbumin-positive basket cells differentiate among hippocampal pyramidal cells. Neuron. 2014; 82(5):1129–1144. doi: http://dx.doi.org/10.1016/j.neuron.2014.03.034.

Lee WCA, Bonin V, Reed M, Graham BJ, Hood G, Glattfelder K, Reid RC. Anatomy and function of an excitatory network in the visual cortex. Nature. 2016; 532(7599):370–374. doi: http://dx.doi.org/10.1038/nature17192.

Letzkus JJ, Wolff SBE, Meyer EMM, Tovote P, Courtin J, Herry C, Lüthi A. A disinhibitory microcircuit for associative fear learning in the auditory cortex. Nature. 2011; 480(7377):331–335. doi: https://doi.org/10.1038/nature10674.

Levakova M, Kostal L, Monsempès C, Lucas P, Kobayashi R. Adaptive integrate-and-fire model reproduces the dynamics of olfactory receptor neuron responses in a moth. Journal of The Royal Society Interface. 2019; 16:20190246. doi: http://dx.doi.org/10.1098/rsif.2019.0246.

Litwin-Kumar A, Doiron B. Formation and maintenance of neuronal assemblies through synaptic plasticity. Nature Communications. 2014; 5(5319). doi: http://dx.doi.org/10.1038/ncomms6319.

Lundstrom BN, Fairhall AL, Maravall M. Multiple timescale encoding of slowly varying whisker stimulus envelope in cortical and thalamic neurons in vivo. Journal of Neuroscience. 2010; 30(14):5071–5077. doi: https://doi.org/10.1523/JNEUROSCI.2193-09.2010.

Ma WP, Liu BH, Li YT, Huang ZJ, Zhang LI, Tao HW. Visual representations by cortical somatostatin inhibitory neurons - Selective but with weak and delayed responses. Journal of Neuroscience. 2010; 30(43):14371–14379. doi: https://doi.org/10.1523/JNEUROSCI.3248-10.2010.

Mackwood O, Naumann LB, Sprekeler H. Learning excitatory-inhibitory neuronal assemblies in recurrent networks. bioRxiv. 2020; 016352. doi: https://doi.org/10.1101/2020.03.30.016352.

Maffei A, Nataraj K, Nelson SB, Turrigiano GG. Potentiation of cortical inhibition by visual deprivation. Nature. 2006; 443(7107):81–84. doi: https://doi.org/10.1038/nature05079.

Makino H, Komiyama T. Learning enhances the relative impact of top-down processing in the visual cortex. Nature Neuroscience. 2015; 18(8):1116–1122. doi: https://doi.org/10.1038/nn.4061.

Mill R, Coath M, Wennekers T, Denham SL. A neurocomputational model of stimulus-specific adaptation to oddball and markov sequences. PLoS Computational Biology. 2011; 7(8):e1002117. doi: https://doi.org/10.1371/journal.pcbi.1002117.

Mill R, Coath M, Wennekers T, Denham SL. Abstract stimulus-specific adaptation models. Neural Computation. 2011; 23:435–476. doi: https://doi.org/10.1162/NECO{\_}a{\_}00077.

Mill R, Coath M, Wennekers T, Denham SL. Characterising stimulus-specific adaptation using a multi-layer field model. Brain Research. 2012; 1434:178–188. doi: http://dx.doi.org/10.1016/j.brainres.2011.08.063.

Miller JEK, Ayzenshtat I, Carrillo-Reid L, Yuste R. Visual stimuli recruit intrinsically generated cortical ensembles. Proceedings of the National Academy of Sciences of the United States of America. 2014; 111:E4053–E4061. doi: https://doi.org/10.1073/pnas.1406077111.

Movshon JA, Lennie P. Pattern-selective adaptation in visual cortical neurones. Nature. 1979; 278(5707):850–852. doi: https://doi.org/10.1038/278850a0.

Näätänen R, Paavilainen P, Rinne T, Alho K. The mismatch negativity (MMN) in basic research of central auditory processing: A review. Clinical Neurophysiology. 2007; 118:2544–2590. doi: https://doi.org/10.1016/j.clinph.2007.04.026.

Näätänen R, Simpson M, Loveless NE. Stimulus deviance and evoked potentials. Biological Psychology. 1982; 14:53–98. doi: https://doi.org/10.1016/0301-0511(82)90017-5.

Najafi F, Elsayed GF, Cao R, Pnevmatikakis E, Latham PE, Cunningham J, Churchland AK. Excitatory and inhibitory subnetworks are equally selective during decision-making and emerge simultaneously during learning. Neuron. 2020; 105(1):165–179. doi: https://doi.org/10.1016/j.neuron.2019.09.045.

Natan RG, Briguglio JJ, Mwilambwe-Tshilobo L, Jones SI, Aizenberg M, Goldberg EM, Geffen MN. Complementary control of sensory adaptation by two types of cortical interneurons. eLife. 2015; 4:e09868. doi: https://doi.org/10.7554/eLife.09868.

Natan RG, Rao W, Geffen MN. Cortical Interneurons Differentially Shape Frequency Tuning following Adaptation. Cell Reports. 2017; 21(4):878–890. doi: https://doi.org/10.1016/j.celrep.2017.10.012.

Nelken I. Stimulus-specific adaptation and deviance detection in the auditory system: experiments and models. Biological Cybernetics. 2014; 108:655–663. doi: https://doi.org/10.1007/s00422-014-0585-7.

Ohki K, Reid RC. Specificity and randomness in the visual cortex. Current Opinion in Neurobiology. 2007; 17:401–407. doi: https://doi.org/10.1016/j.conb.2007.07.007.

Park Y, Geffen MN. A circuit model of auditory cortex. PLoS Computational Biology. 2020; 15(7):e1008016. doi: http://dx.doi.org/10.1371/journal.pcbi.1008016.

Pfister JP, Gerstner W. Triplets of Spikes in a Model of Spike Timing-Dependent Plasticity. Journal of Neuroscience. 2006; 26(38):9673–9682. doi: https://doi.org/10.1523/jneurosci.1425-06.2006.

Ramaswami M. Network plasticity in adaptive filtering and behavioral habituation. Neuron. 2014; 82(6):1216–1229. doi: http://dx.doi.org/10.1016/j.neuron.2014.04.035.

Rao RPN, Ballard DH. Predictive coding in the visual cortex: a functional interpretation of some extra-classical receptive-field effects. Nature Neuroscience. 1999 1; 2(1):79–87. doi: https://doi.org/10.1038/4580.

Ross JM, Hamm JP. Cortical Microcircuit Mechanisms of Mismatch Negativity and Its Underlying Subcomponents. Frontiers in Neural Circuits. 2020; 14(13). doi: https://doi.org/10.3389/fncir.2020.00013.

Rost T, Deger M, Nawrot MP. Winnerless competition in clustered balanced networks: inhibitory assemblies do the trick. Biological Cybernetics. 2018; 112:81–98. doi: https://doi.org/10.1007/s00422-017-0737-7.

Rostami V, Rost T, Riehle A, Albada SJv, Nawrot MP. Spiking neural network model of motor cortex with joint excitatory and inhibitory clusters reflects task uncertainty, reaction times, and variability dynamics. bioRxiv. 2020; 968339. doi: https://doi.org/10.1101/2020.02.27.968339.

Schwartz G, Berry II MJ. Sophisticated temporal pattern recognition in retinal ganglion cells. Journal of Neurophysiology. 2008; 99:1787–1798. doi: https://doi.org/10.1152/jn.01025.2007.

Schwartz G, Harris R, Shrom D, Berry II MJ. Detection and prediction of periodic patterns by the retina. Nature Neuroscience. 2007 5; 10(5):552–554. doi: https://doi.org/10.1038/nn1887.

Simoncelli EP, Olshausen BA. Natural Image Statistics and Neural Representation. Annual Review of Neuroscience. 2001; 24:11931216. doi: https://doi.org/10.1146/annurev.neuro.24.1.1193.

Sjöström PJ, Turrigiano GG, Nelson SB. Rate, Timing, and Cooperativity Jointly Determine Cortical Synaptic Plasticity. Neuron. 2001; 32(6):1149–1164. doi: https://doi.org/10.1016/s0896-6273(01)00542-6.

Snow M, Coen-Cagli R, Schwartz O. Adaptation in the visual cortex: a case for probing neuronal populations with natural stimuli. F1000Research. 2017; 6(F1000 Faculty Rev):1246. doi: https://doi.org/10.12688/f1000research.11154.1.

Spratling MW. A review of predictive coding algorithms. Brain and Cognition. 2017; 112:92–97. doi: http://dx.doi.org/10.1016/j.bandc.2015.11.003.

Sussman ES, Chen S, Sussman-Fort J, Dinces E. The five myths of MMN: Redefining how to Use MMN in basic and clinical research. Brain Topography. 2014; 27:553–564. doi: https://doi.org/10.1007/s10548-013-0326-6.

Taaseh N, Yaron A, Nelken I. Stimulus-specific adaptation and deviance detection in the rat auditory cortex. PLoS ONE. 2011; 6(8):e23369. doi: https://doi.org/10.1371/journal.pone.0023369.

Thompson A, Gribizis A, Chen C, Crair MC. Activity-dependent development of visual receptive fields. Current Opinion in Neurobiology. 2017; 42:136–143. doi: http://dx.doi.org/10.1016/j.conb.2016.12.007.

Tikhonravov D, Neuvonen T, Pertovaara A, Savioja K, Ruusuvirta T, Näätänen R, Carlson S. Effects of an NMDA-receptor antagonist MK-801 on an MMN-like response recorded in anesthetized rats. Brain Research. 2008; 1203:97–102. doi: https://doi.org/10.1016/j.brainres.2008.02.006.

Tremblay R, Lee S, Rudy B. GABAergic Interneurons in the Neocortex: From Cellular Properties to Circuits. Neuron. 2016; 91(2):260–292. doi: https://doi.org/10.1016/j.neuron.2016.06.033.

Tsodyks M, Pawelzik K, Markram H. Neural Networks with Dynamic Synapses. Neural Computation. 1998; 10:821–835. doi: https://doi.org/10.1162/089976698300017502.

Ulanovsky N, Las L, Farkas D, Nelken I. Multiple time scales of adaptation in auditory cortex neurons. Journal of Neuroscience. 2004; 24(46):10440–10453. doi: https://doi.org/10.1523/JNEUROSCI.1905-04.2004.

Ulanovsky N, Las L, Nelken I. Processing of low-probability sounds by cortical neurons. Nature Neuroscience. 2003; 6(4):391–398. doi: https://doi.org/10.1038/nn1032.

Vinken K, Vogels R, Op de Beeck H. Recent Visual Experience Shapes Visual Processing in Rats through Stimulus-Specific Adaptation and Response Enhancement. Current Biology. 2017 3; 27(6):914–919. doi: http://dx.doi.org/10.1016/j.cub.2017.02.024.

Vogels TP, Sprekeler H, Zenke F, Clopath C, Gerstner W. Inhibitory Plasticity Balances Excitation and Inhibition in Sensory Pathways and Memory Networks. Science. 2011; 334(6062):1569–1573. doi: https://doi.org/10.1126/science.1211095.

Wang L, Maffei A. Inhibitory Plasticity Dictates the Sign of Plasticity at Excitatory Synapses. Journal of Neuroscience. 2014; 34(4):1083–1093. doi: https://doi.org/10.1523/jneurosci.4711-13.2014.

Wang XJ, Yang GR. A disinhibitory circuit motif and flexible information routing in the brain. Current Opinion in Neurobiology. 2018; 49:75–83. doi: http://dx.doi.org/10.1016/j.conb.2018.01.002.

Weber AI, Krishnamurthy K, Fairhall AL. Coding Principles in Adaptation. Annual Review of Vision Science. 2019; 5:427–449. doi: https://doi.org/10.1146/annurev-vision-091718-014818.

Woodin MA, Ganguly K, Poo MM. Coincident Pre-and Postsynaptic Activity Modifies GABAergic Synapses by Postsynaptic Changes in Cl-Transporter Activity. Neuron. 2003; 39(5):807–820. doi: https://doi.org/10.1016/s0896-6273(03)00507-5.

Xue M, Atallah BV, Scanziani M. Equalizing excitation-inhibition ratios across visual cortical neurons. Nature. 2014; 511:596–600. doi: https://doi.org/10.1038/nature13321.

Yarden TS, Nelken I. Stimulus-specific adaptation in a recurrent network model of primary auditory cortex. PLoS Computational Biology. 2017; 13(3):e1005437. doi: https://doi.org/10.1371/journal.pcbi.1005437.

Yaron A, Hershenhoren I, Nelken I. Sensitivity to Complex Statistical Regularities in Rat Auditory Cortex. Neuron. 2012; 76(3):603–615. doi: http://dx.doi.org/10.1016/j.neuron.2012.08.025.

Yaron A, Jankowski MM, Badrieh R, Nelken I. Stimulus-specific adaptation to behaviorally-relevant sounds in awake rats. PLoS ONE. 2020; 15(3):e0221541. doi: https://doi.org/10.1371/journal.pone.0221541.

Zenke F, Agnes EJ, Gerstner W. Diverse synaptic plasticity mechanisms orchestrated to form and retrieve memories in spiking neural networks. Nature Communications. 2015; 6(6922). doi: https://doi.org/10.1038/ncomms7922.

Zmarz P, Keller GB. Mismatch Receptive Fields in Mouse Visual Cortex. Neuron. 2016; 92(4):766–772. doi: http://dx.doi.org/10.1016/j.neuron.2016.09.057.

Znamenskiy P, Kim MH, Muir DR, Iacaruso F, Hofer SB, Mrsic-Flogel TD. Functional selectivity and specific connectivity of inhibitory neurons in primary visual cortex. bioRxiv. 2018; 294835. doi: http://dx.doi.org/10.1101/294835.

Zucker RS, Regehr WG. Short-term synaptic plasticity. Annual Review of Physiology. 2002; 64:355–405. doi: https://doi.org/10.1146/annurev.physiol.64.092501.114547.

